# Comparative Computational Modeling of Approach–Avoidance Biases in Suicidal Populations via Hierarchical Bayesian Inference

**DOI:** 10.1101/2025.08.26.672271

**Authors:** Pamina Laessing, Povilas Karvelis, James Kennedy, Clement C. Zai, Peter Dayan, Andreea Diaconescu

## Abstract

Pavlovian “approach or avoid” impulses are critical behavioral biases that, in excess, are linked to multiple psychiatric conditions. To investigate how such biases contribute to suicidal thoughts and behaviors, we analyzed data from two clinical populations completing an aversive Go/NoGo task. This task disentangles motor action (Go or NoGo) from outcome valence (escape from, or avoidance of, an aversive stimulus), enabling the isolation of Pavlovian biases from instrumental learning processes.

We compared multiple computational models that had previously been proposed to explain Pavlovian tendencies, including reinforcement learning, active inference, and drift diffusion–based approaches. We employed a hierarchical Bayesian inference procedure that treats model identity as a random factor at the individual level, allowing an unbiased determination of which mechanisms most accurately captured participants’ behavior. Across both datasets, models featuring Pavlovian context biases plus a value-decay mechanism best accounted for performance. By contrast, policy-based Pavlovian models and more complex approaches, such as those integrating working memory or active inference, were supported by fewer study participants.

These findings suggest that reflexive biases exert a persistent influence on decision-making, and that value decay plays a critical role in shaping behavior over time. Our results demonstrate the importance of systematically comparing and accounting for relevant cognitive processes to explain observed task behaviors. Understanding the factors contributing to task performance may help clarify how Pavlovian tendencies relate to psychopathology, including, in our case, elevated suicide risk. Finally, we illustrate how a complete hierarchical model selection framework can be applied to identify the most plausible mechanisms underlying Pavlovian biases, offering a robust approach for advancing our understanding of task behaviors and establishing clinical utility in future studies.

**Author summary:** Automatic “approach or avoid” reactions shape behavior, particularly in stressful or negative situations. In this study, we explored how these reflex-like tendencies might contribute to suicidal thoughts and behaviors. Two clinical groups completed a computerized task measuring responses to unpleasant sounds. Participants made either active responses (pressing a button to stop a sound) or passive responses (refraining from pressing to avoid starting a sound), allowing us to examine the interplay of automatic impulses and learning from past experiences.

Our analysis showed that behavior was best explained by a model combining stable “approach or avoid” impulses with a forgetting process that reduced reliance on past experiences over time. More complex models involving strategies or memory-based control were less effective.

These findings suggest that individuals with suicidal tendencies may rely on persistent reflex-like behaviors and over-index recent outcomes, compromising their ability to learn in uncertain environmental conditions. Understanding these cognitive processes provides insights into why some individuals feel trapped in harmful patterns of thought and behavior. Our work highlights how identifying shared traits in clinical populations using model-based methods can inform targeted mental health interventions and improve our understanding of cognitive functioning across disorders.

## Introduction

A Pavlovian system of control orchestrates default, reflexive approach and avoidance behaviors in response to predictions of reward and punishment (Dickinson, 1980; Mackintosh, 1983). It is complemented by an instrumental system which learns, typically more slowly, to perform actions contingent on their outcomes. The integration of these control systems is crucial for achieving adaptive behavior, striking a balance between faster, reflexive responses and the flexibility afforded by learned actions (N. D. Daw et al., 2005).

While the default actions provided by the Pavlovian system have likely been evolutionarily advantageous, problems ensue when Pavlovian behaviors are inconsistent with instrumental requirements [1–3]. Indeed, excess influence of Pavlovian biases have been associated with several psychiatric disorders, including, but not limited to, PTSD [4], mood and anxiety disorders [5], addiction disorders [6] and suicidality [7].

The transdiagnostic prevalence of maladaptive Pavlovian tendencies underscores the importance of understanding the mechanisms that underlie them. Computational psychiatry employs reward/punishment tasks - such as the orthogonalized Go/NoGo task [8, 9] - that disentangle reflexive Pavlovian biases from other cognitive processes, providing a window into these mechanisms. However, with just a couple of notable exceptions [7, 10], such Go/NoGo tasks have been applied in a monetary context, even though, for many psychiatric conditions, negative contingencies involving avoidance of, or escape from, a primary aversive stimulus, would be more obviously relevant. Here, we sought to use suicidality [7] as a critical test case for elucidating Pavlovian-instrumental competition.

In computational psychiatry, it is conventional to proceed by comparing the fit and the inferred parameter values of computational models that incorporate and differentiate separate possible mechanisms. Although the orthogonalized Go/NoGo task primarily reflects Pavlovian tendencies, other facets such as valence, reward/punishment stimuli, task length, and reversals—can interact with model-free and model-based learning systems in the context of instrumental learning, working memory, and other competing processes [11, 12]. Thus, here, we perform a systematic, head-to-head, comparison not only within, but also between, the two main modelling frameworks that have been suggested: reinforcement learning (e.g., [9–11, 13, 14] and active inference [15, 16]. We do this using behavioral data from two clinical populations with suicidal thoughts and behaviors (STBs). Our analysis reveals how task performance can be understood through distinct cognitive mechanisms, offering valuable insights, while also surfacing inherent limitations in the data and methodologies.

### Assessing Pavlovian tendencies in suicidality

Suicidality is a complex phenomenon with a highly heterogeneous clinical expression and poorly understood cognitive and neurobiological risk factors. Although it is traditionally defined as a uniform concept, the presentation of suicidal ideation and behaviours varies across psychopathologies, biological predispositions, and psychosocial contexts (see [17] for an overview), leading to an array of associated cognitive deficits. Thus, in recent years, stratification approaches have been used to identify subgroups of suicidality based on personality traits, clinical comorbidities, and temporal patterns. [17–21]. However, small sample sizes, inconsistent definitions and findings, and limited clinical data collection have hindered the development of a consensus on meaningful subgroup identification.

One promising approach stems from the observation that STB is associated with deficits in reinforcement learning (RL) tasks. These deficits include impairments in active-escape behaviors in aversive contexts [10], probabilistic reversal learning [22], and value-based decision-making [23]. Such impairments appear to vary in severity across subgroups and may reflect an increased influence of Pavlovian over instrumental control, particularly in negatively valenced social contexts [24]. Earlier studies have explored this by quantifying and modelling performance biases in tasks in which Pavlovian and instrumental contingencies are congruent vs incongruent [10, 23].

In this study, we adopt an empirical model-driven approach to investigate STB and the potential heterogeneity of associated behavior. We analyze and model behavior in negatively valenced orthogonalized Go/NoGo tasks [7, 9, 10] and compare computational models of learning and decision-making mechanisms that can generate Pavlovian biases using an empirical hierarchical Bayesian inference approach, based on the method outlined in [25]. This framework treats model identity as a random effect at the individual level in the model fitting process, enabling unbiased assessment of computational mechanisms at both individual and group levels. By leveraging this robust statistical approach, we provide detailed insights into the cognitive processes associated with STB.

## Materials and Methods

To examine the cognitive mechanisms underlying STB, we analyzed two datasets involving aversive Go/NoGo tasks designed to measure reflexive and learned responses to negative stimuli (Figure 1). These complementary datasets allowed us to apply computational modeling and hierarchical Bayesian inference to identify the mechanisms driving observed behaviors and evaluate hypotheses about Pavlovian biases in decision-making.

**Fig 1.**
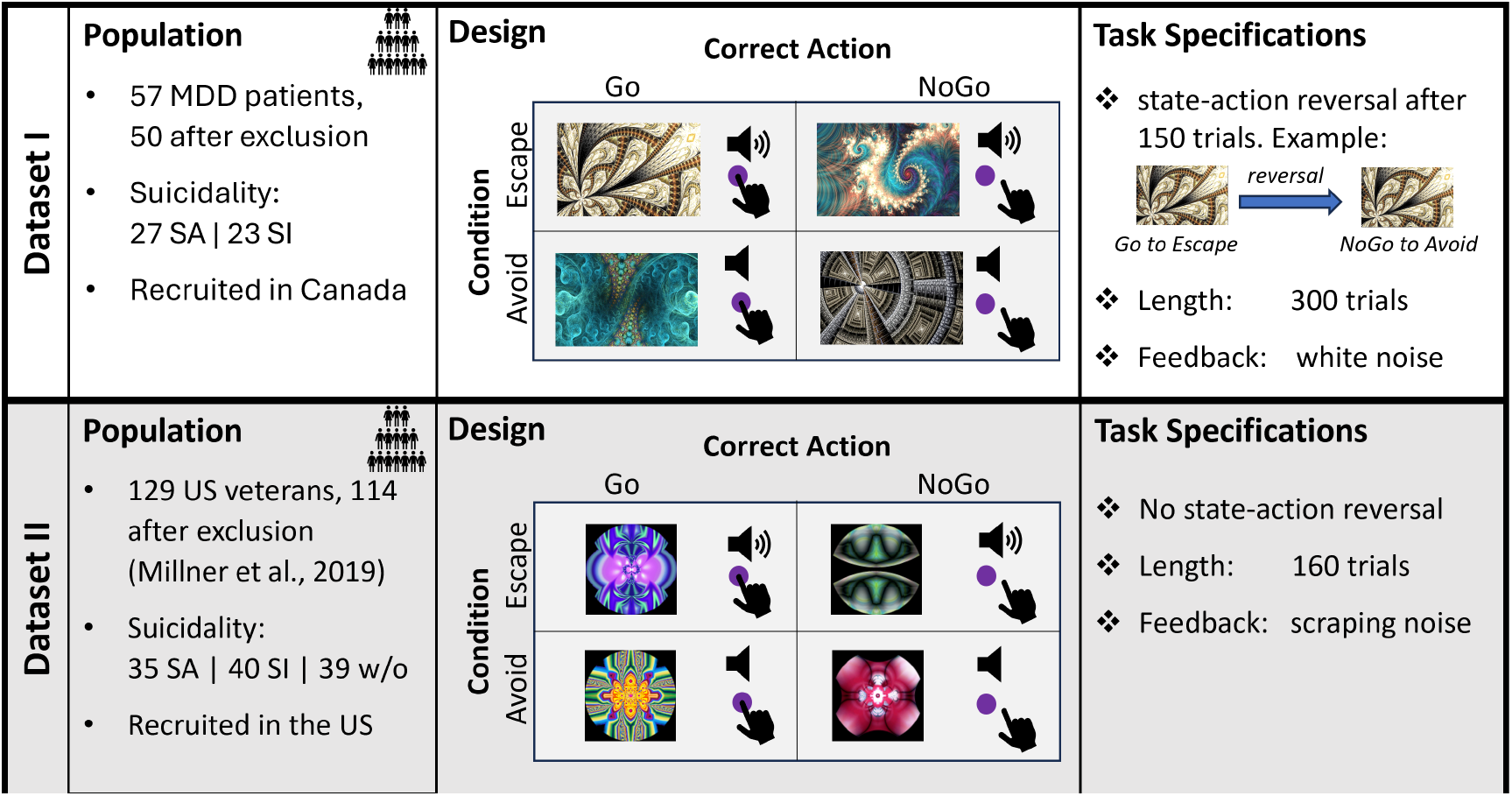
Overview of the datasets. The computational model space was explored on two datasets collected in distinct populations. The aversive Go/NoGo design employed in each population differed in design and length. *Abbreviations: MDD-major depressive disorder, SI-Suicidal Ideation, SA-Suicidal Attempt

### Participants

#### Dataset I

Participants were drawn from a clinical study aimed at identifying suicidal subgroups based on behavioral, psychological, and genomic factors. We analyzed an early subsample of this study. Participants were re-contacted from IMPACT (Individualized Medicine: Pharmacogenetic Assessment and Clinical Treatment), an ongoing provincially funded pharmacogenetics study [26]. The cohort included adult participants with existing diagnoses of anxiety, depression, psychosis, pain, ADHD, bipolar disorder, tic disorders, or sleep disorders. Participants received $60 for completing the study assessments, including the Go/NoGo task. Of the 57 participants who completed the task, 50 (72% female, mean age: 39.6±12.4) were included after exclusion based on outlier performance measures. (see SI)

The study included a comprehensive assessment battery. Suicidal behavior was characterized using the Columbia-Suicide Severity Rating Scale (CSSR-S, [27]), and additional self-report scales evaluated depression, hopelessness, and anxiety. Of the analyzed participants, 27 had experienced suicidal ideation (SI), and 23 had attempted suicide (SA), with 3 participants reporting recent SA. The online Go/NoGo task included 300 trials, and behavioral outcomes such as choice and reaction time were recorded.

#### Dataset II

This dataset was reanalyzed from a study by [10], who recruited 158 veterans from Boston-area treatment centers. Of these, 127 participants were psychiatric inpatients, and 31 were outpatients. The original study published data from 129 participants after excluding the first participants due to a change in the task presentation. After applying exclusion criteria consistent with Dataset I, 114 participants (28% female, mean age: 40.6±13.5) were included in our analysis: 40 with a history of SI , 35 with a history of SA, and 39 with a history of psychiatric conditions other than STB. Participants completed an in-person aversive Go/NoGo task with 160 trials. Behavioral outcomes, including choice and reaction time, were recorded. Psychological assessments included, among others, the Self-Injurious Thoughts and Behaviors Interview (SITBI) [28] for validity, the 9-item Patient Health Questionnaire (PHQ-9; [29]) for depressive symptoms, the Beck Hopelessness Scale (BHS; [30]) and the UPPS Impulsive Behavior Scale (UPPS; [31]). Further details about the dataset and assessments can be found in the original publication [10].

### Ethics Statement

The IMPACT study (Dataset I) was approved by the Centre for Addiction and Mental Health Research Ethics Board - approval: 021/2016-03. All participants provided written informed consent to participate in this study.

Dataset II was reanalyzed from a previous study, approved by the VA Boston Healthcare System (Institutional Review Board [IRB] 2773) and Harvard University (IRB 13-1390) IRBs. Further details can be found in the original publication [10].

### Cognitive Task

The study employs orthogonal Go/NoGo tasks to assess Pavlovian tendencies in the context of suicidality. The Go/No-Go task is a widely used paradigm in behavioural psychology to elucidate the mechanisms underlying Pavlovian biases in decision-making [9, 32–34]. These tasks require participants to learn to perform or withhold actions (Go or NoGo) in response to specific cues to optimize outcomes, with auditory and visual feedback provided based on their responses. Participants’ primary goal was to maximize neutral feedback (silence) by selecting the appropriate response to each cue. The tasks include both escape and avoid conditions: in the escape condition, participants terminate an aversive stimulus, while in the avoid condition, they prevent its onset. Four fractal image cues were used, each corresponding to one of four conditions: Go-to-Escape, Go-to-Avoid, NoGo-to-Escape, and NoGo-to-Avoid. Feedback contingencies were probabilistic, with optimal responses resulting in silence 80% of the time, the remaining 20% resulting in the aversive stimulus. Suboptimal responses reversed these probabilities.

Two versions of the Go/NoGo task were utilized, differing in the number of trials, feedback stimuli, and task structure.

In Dataset I, the task comprised 300 trials, with a mid-task cue-action-outcome reversal after 150 trials to increase complexity. Aversive feedback consisted of white noise. Participants adjusted headphone volume to an unpleasant but tolerable level. Subjective experience probes, including task difficulty and controllability, were included. Trial types were chosen uniformly at random, with each being presented approximately 75 times, albeit slightly unevenly due to the reversal (e.g., NGA:67, GA:79, NGE:78, GE:76).

In Dataset II, the task included 160 trials with no cue-action-outcome reversal. Neutral feedback was silence, while aversive feedback consisted of a scraping sound at 80–85 dB. Feedback duration was standardized to align trial lengths across conditions. Each cue appeared in 40 trials in permuted order.

### Data Analyses

#### Behavioural Data

Behavioural data were initially analysed using both model-agnostic metrics and simple model-based heuristics (Win-Stay and Lose-Switch strategies). We computed accuracy (ratio of trials with correct responses to all trials) on the overall task, as well as for each of the four cue types, and before and after the reversal for dataset I. A behavioural Go bias was defined as the difference in accuracy for Go trials (GA and GE) relative to NoGo trials (NGA and NGE). Similarly, a Go-to-Escape bias was calculated as the accuracy difference between Go and NoGo trials in the Escape condition, while a NoGo-to-Avoid bias was the accuracy ratio between NoGo and Go trials in the Avoid condition.

#### Modelling Pavlovian Biases (Model Space)

We built and tested a range of computational models that are commonly applied to the Go/NoGo task. The models formalize and combine aspects of learning and Pavlovian mechanisms in distinct ways, offering insights into the variability observed in behaviour.

The nomenclature used in the following sections is modular. If a mechanism is not used in the model, we indicated its absence with the *-symbol 2. The nomenclature is as follows:

1. Learning Mechanism: R- Reinforcement Learning, K- Kalman filter, A- Active Inference
  a. Pavlovian Mechanism: b- context bias, a-evidence bias, *ω*-policy, *ν*-policy with variable contribution
  b. Go-bias: G-global Go bias
  c. Learning: *ρ* - reward sensitivity by condition
  d. Mnemonic mechanism: f-forgetting, m-working memory

Below, we introduce in further detail the modeling mechanisms considered in this work.

### Model Families: Reinforcement Learning and Active Inference

We evaluated models from two primary families: Reinforcement Learning (RL) and Active Inference (AI). These families capture different facets of decision-making:

I. **Active Inference (A):** AI agents minimize surprise by aligning predicted outcomes with prior preferences, performing inference to optimize behaviour rather than directly maximizing reward.
II. **Reinforcement Learning (R & K):** Model-free RL agents maximize reward by learning state-action values and selecting actions based on accumulated experience.

Mechanisms embedded in both families address Pavlovian biases, learning strategies, reward sensitivity, action selection, and recency effects.

### Active Inference Models

Developed by Karvelis & Diaconescu [16], the **Active Inference Suicide Model** (*A*_*ωG*\*\*_) parametrizes hopelessness and Active Escape biases in suicidal populations. Pavlovian tendencies are framed as a fixed policy which exists alongside instrumental Go/NoGo action policies. The Pavlovian policy shapes expectations, influencing state-action selection to align with prior beliefs.

The model is parametrized as follows. Parameter ‘c’ encodes the prior dis-preference for states. Here, we assume equal preference against all states with an aversive sound compared to ‘neutral’ states, reducing the parameter to a single fitted value. *z*_*E*_ and *z*_*A*_ represent the belief that Pavlovian actions (Go-to-escape and NoGo-to-avoid) lead to preferred outcome states. Pavlovian policies are fixed over trials for this model. *d*_*m*_ is a belief decay threshold which defines the threshold, above which state action prediction errors (SAPEs) lead to significant belief decay in instrumental policies. The model dynamically adjusts learning rates through state-action-prediction errors (SAPEs). Learning extends to counterfactual updates and world-model adjustments via state-transition matrices.

Action probabilities are computed using a SoftMax function, reflecting the likelihood that an action results in the expected outcome. *α*_*prec*_ is the temperature parameter defining the slope of the SoftMax function for action selection. We made slight adjustments to the original *A*_*ωG*\*\*_ model to accommodate the aversively balanced orthogonal Go/NoGo task by incorporating a prior preference for Go policies *p*_*Go*_, to accommodate the general action biases observed in participants. For the computational details of the model, we refer the interested reader to the original publication [16].

### Reinforcement Learning Models

Most RL models considered were based on the Rescorla-Wagner or delta-rule framework [35], which is known for its simplicity and efficiency in modelling associative learning. The model maintains state-action values, or Q-values, which are updated based on the reward r and learning rate *lr* after each trial:

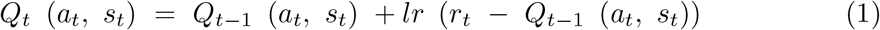

Following extensive previous work on versions of the orthogonal Go-NoGo task and models [7, 9, 10, 32], we consider modelling mechanisms which specifically capture Pavlovian tendencies, the influence of working memory and different learning and choice mechanisms, as well as their interplay.

#### 1. Pavlovian Tendencies

Pavlovian tendencies may contribute to value formation in two major ways:

I. **Pavlovian Context Parameters:** Pavlovian tendencies can be modelled as fixed biases that persist throughout the task [15, 36]. As instrumental learning progresses and state-action values grow, the influence of these biases gradually diminishes. Biases are added to the computed Q-values to form a final state-action value W for action taking:

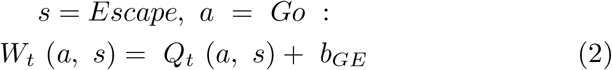

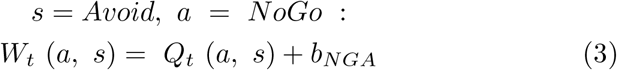
II. **Pavlovian Policies:** Here, the inherent strength of the Pavlovian tendencies is proportional to the values of the states concerned and so remains malleable during the task. The way these Pavlovian policies are combined with instrumental preferences can either be with a fixed contribution *ω* or a variable contribution based on the posterior likelihood of the environment being uncontrollable (i.e. Pavlovian), adapting to task conditions over time [13]. Our models assumed equal learning rates for Pavlovian and instrumental policies, consistent with prior research suggesting overlap in their underlying mechanisms [12]. Pavlovian Policies are modelled as arising from action-independent state-values *V* that are learned as:

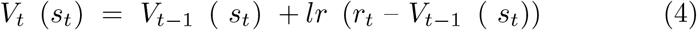

Instrumental and Pavlovian policies are combined with a mixing parameter *ω* to obtain a final state-action propensity *W* for action-taking:

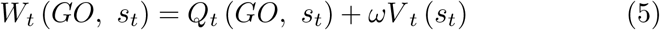

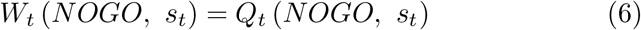
III. **Pavlovian Evidence Bias**: Based on models of the evidence accumulation that leads an agent to take a decision, Pavlovian tendencies can be embedded as biased starting points in the accumulation process [7, 10]. Reinforcement Learning-Drift Diffusion models (RL-DDM; [37, 38]) extend classical RL models by capturing the translation of state-action values into motor responses. Decision-making is represented as a noisy evidence accumulation process [38] that allows the modelling of reaction time. Introducing Pavlovian biased starting points into the evidence accumulation process biases the decision-making process. Overcoming it requires strong instrumental values, reinforcing Pavlovian-congruent actions in the early stages of the task. While RL-DDM models typically capture both choice and reaction time data, simplified versions were employed here to model choice data exclusively. This required reducing the parameter space to ensure parameter recoverability. The simplified RL-DDM models employed here (*R*_*α*\*\*\*_ and *R*_*α*\**ρ*\*_) translate action values into drift velocity *v*

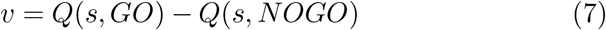

This drift velocity is then fed into the Drift Diffusion model, where the probability for a Go/NoGo action is defined as the probability of reaching the Go/NoGo limit first, given the velocity and starting point. The starting points *z*_*A*_ for avoid conditions and *z*_*E*_ for escape conditions are defined as

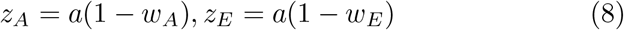

where *a* is the boundary separation between the NoGo and Go choice boundaries and *w*_*A*_ and *w*_*E*_ are the (normalized) avoid and escape biases. The probability of the first passage time at the upper choice boundary *c* is calculated according to [39].

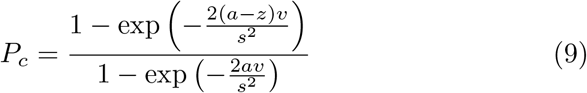

where *s* is a scaling factor set to 1 for modelling. In this model of Pavlovian tendencies, the bias shapes the SoftMax decision function through a combination of shift along the x-axis and adjustment of the slope (see Figure S1 for further details). Mechanically, this means that stronger Pavlovian biases increase the threshold and decrease the sensitivity for Pavlovian-incongruent instrumental values to significantly influence action taking. Conversely, Pavlovian context bias parameters only effect the threshold by shifting the SoftMax function along the x-axis, but not the sensitivity, proposing a clearer separation between the effects of Pavlovian and instrumental learning systems on action-taking. (see SI 7.2.1 for illustrative examples)

#### 2. Global Action (Go) Bias

A Go-bias parameter captures global, context-independent tendencies towards action taking, operationalized here as Go responses. For RL models, a bias parameter *b* is added to the action weights of Go responses. For most models considered here, a separate Go bias parameter is not recoverable given the studied datasets. We only included the parameter for models in which it was recoverable (ICC*>*.4 for all parameters), here the RL-policy, *R*_*νG*\*\*_ and *R*_*ωG*\*\*_ models.

#### 3. Learning Mechanisms

To account for potential differences in reinforcement sensitivity in different environmental contexts, we introduce separate values *ρ*_*A/E*_ for Avoid and Escape conditions. The new update equations are:

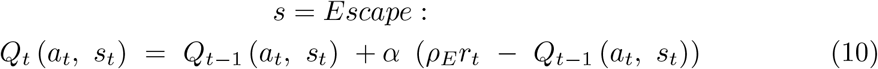

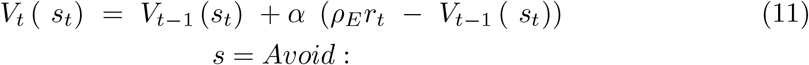

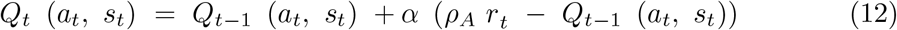

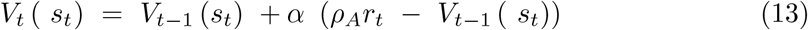

The *A*_*ωG*\*\*_ model boasts a dynamic learning rate based on state-prediction errors. By contrast, Rescorla-Wagner models typically employ fixed learning rates. To create a dynamic learning rate in our RL models, we consider a simple Kalman filter (*K*_*b*\*\*\*_), presenting a Bayesian generalization of Rescorla-Wagner [40–42]. In this model, the agent tracks a posterior distribution instead of a point estimate of the outcome, utilizing the uncertainty to dynamically adjust its learning rate to the environmental volatility. The Kalman gain replaces the learning rate *α* in the Rescorla-Wagner model.

Importantly, the Kalman gain is stimulus-specific and changes dynamically with each trial n:

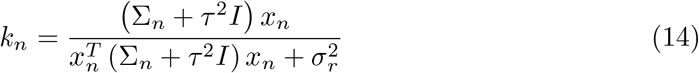

where Σ_*n*_ is the covariance matrix of the posterior distribution over the stimulus-action value, *τ* is the diffusion variance and *σ* is the prior variance.

#### 4. Choice Mechanism

The translation of action values into choice probabilities is based on a simple SoftMax function for all models, except the RL-DDM models (*R*_*a*\*\*\*_ and *R*_*α*\**ρ*\*_), which has its own modified SoftMax function (see equ. 8).

In the simple SoftMax functions, decision noise was modelled by tuning the function using either a temperature parameter *T* (used in *A*_*bG*\*\*_) or its inverse, 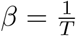

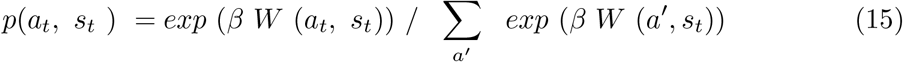

In some of the proposed models, the role of the inverse temperature parameter is taken by the reward sensitivity (*ρ*) parameter. This introduces a condition-dependent sensitivity to action-value differences. In models with reward sensitivity, we additionally considered a lapse parameter *ξ*, allowing the model to account for value-to-choice translation errors and random action lapses. This lapse parameters acts as irreducible noise.

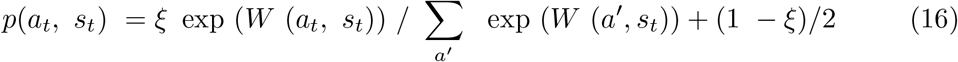

Parametrization choices for all models have been vetted for parameter recoverability (see SI 7.5.2).

#### 5. Mnemonic Mechanisms (R_b**f_ and R_b**m_)

Finally, we consider different mnemonic mechanisms, one proposing uniform forgetting and one with perfect learning, which increases emphasis on the last feedback obtained. The *R*_*b*\*\**f*_ model uses a decay parameter to represent the forgetting of action values over time. The *R*_*b*\*\**m*_ model [11] integrates a working memory with fixed capacity. The working memory system is designed as a Q-learning system with learning rate 1 and a decay factor, capturing degrading memory over time.

These models address task characteristics where inter-trial intervals vary, allowing value decay or memory-driven retention of cue-action mappings over time.

The model space explored here is not exhaustive in terms of general learning mechanisms that may influence task behaviour. We did however explore state-independent learning [43, 44] but did not find evidence justifying the extension of our already large model space for this learning mechanism, as the overall model fit was worse than for state-action learning models. Initially, we also considered a second AI model [15], which incorporates Pavlovian influences as prior policy preferences with a working memory component. However, adapting this model for the Go/NoGo task resulted in poor fitting performance. As such, this model was excluded from further analysis.

### Empirical hierarchical Bayesian inference

Hierarchical Bayesian inference (HBI) is a robust statistical framework that concurrently fits and compares models across individuals and groups [25]. HBI treats model identity as a random effect and uses an expectation-maximization-based iterative estimation approach to update group priors and individual parameter fits. This approach accounts for individual variability while identifying shared group-level patterns in a way that avoids untoward group biases. This makes it particularly suited to provide a nuanced understanding of the underlying cognitive mechanisms in heterogeneous populations, such as those with STB, and is specifically relevant in fields such as psychology and neuroscience, where understanding the interplay between individual differences and group dynamics is essential for developing effective interventions and treatments. For instance, in studies involving psychiatric disorders, HBI can help identify and model subsets of individuals corresponding to disease subgroups. This ability to model heterogeneity not only enhances the accuracy of predictions but also aids in the identification of specific factors that contribute to individual differences in behaviour and cognition.

The original HBI procedure was based on a mean-field approximation with fixed priors. We adapted it to use an empirical variational Bayesian fitting approach with flat priors, equally leveraging individual maximum-a-posteriori (MAP) parameter estimates at the individual level. Both empirical variational Bayesian modeling and mean-field variational Bayes are prominent methods in Bayesian inference, each offering distinct advantages. By deploying flat group priors, the empirical approach lets the data fully shape the posterior distributions, which can improve model performance in data-limited settings and mitigate the need for precisely defined priors [45, 46]. In contrast, mean-field variational Bayes optimization relies on fixed group priors and approximates the posterior distribution by assuming independence among latent variables, thereby simplifying computations but potentially introducing biases if this independence assumption does not hold [47, 48]. While we did not perform formal testing beyond the limited experimental scope here, the adapted empirical HBI (eHBI) procedure showed improved convergence performance for the specific model spaces and datasets tested in this work. These results may be specific to our work and may not generalize to other model spaces and datasets without further investigation.

The final eHBI procedure is depicted in Fig. 2 and combines the original unbiased extended hierarchical fitting from [25] with the improved convergence properties of empirical Bayesian optimization. The main adjustment lies in the second level, where we replaced the Bayesian estimate of group parameters, using a Gaussian-Gamma distribution as prior, with a simpler, empirical estimate of group parameters.

**Fig 2.**
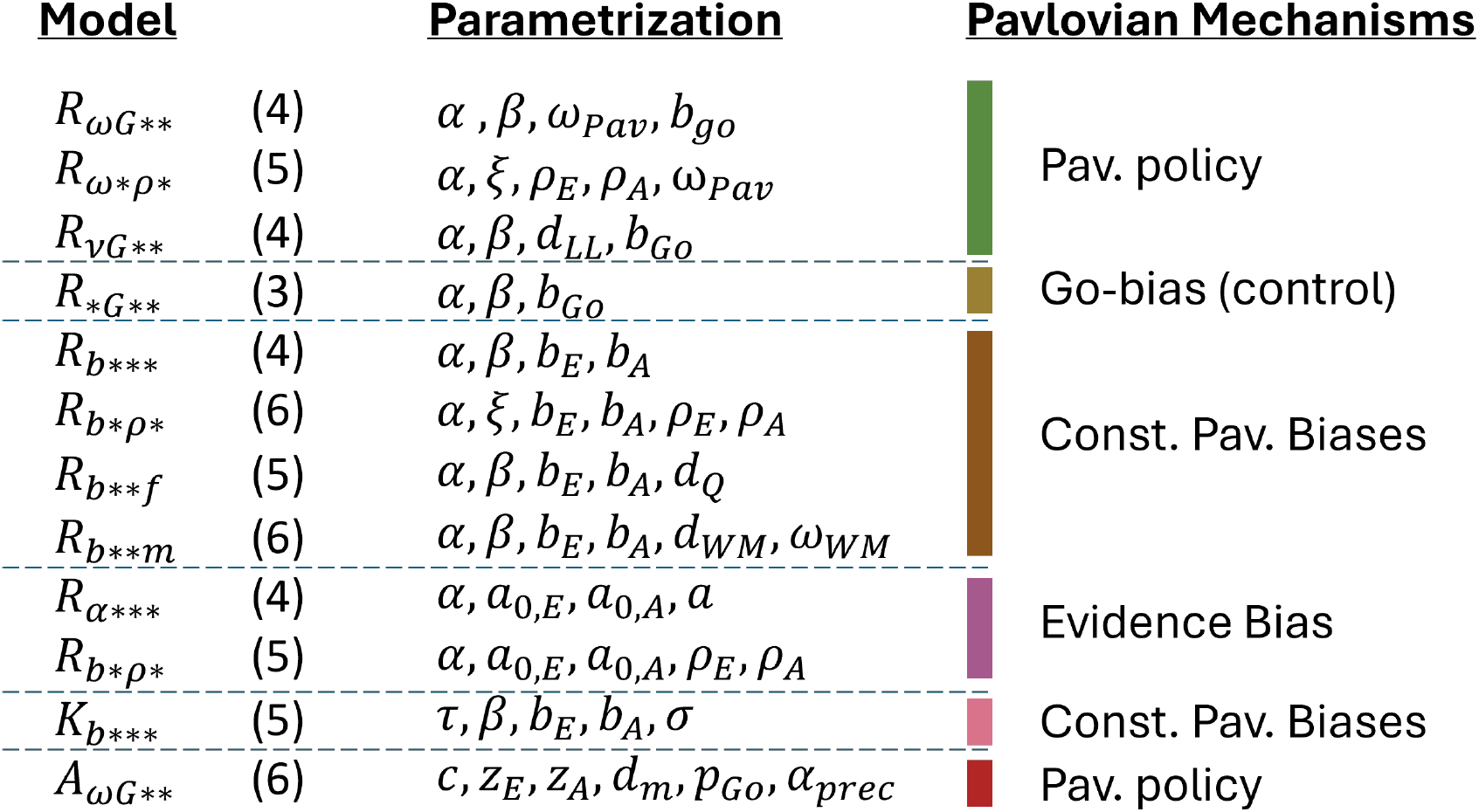
Overview of the full model space for both datasets. Models are categorized based on the model family (Reinforcement Learning -RL, Kalman filter-K or Active Inference-A) and Pavlovian mechanisms, including separate Pavlovian Policy implementations and different instantiations of Pavlovian Bias parameters. Models are parameterized according to these principles. lr is the learning rate, noise is either modeled with β as an inverse temperature or ξ as a decision lapse parameter, ω denotes policy weights, ρ denotes reward sensitivity parameters, d are decay parameters and b are constant bias parameters. The simplified evidence bias models are simplified RL-DDM models, here parametrizing evidence accumulation only with boundary separation a and biased starting points 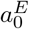 and 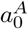 for the evidence accumulation. In the K_b***_ filter, the learning rate is derived from the diffusion rate τ and the state-uncertainty σ (or observation noise). In the Active Inference model A_ωG**_, c is the prior preference over states, z_E_ and z_A_ encode how likely an agent thinks Pavlovian actions lead to the desired outcome, d_m_ is the belief decay threshold, p_Go_ is a prior preference for the Go-policy and T is the action-selection precision (details can be found in [16].

### Model selection procedure

As noted, the eHBI procedure realizes model selection and subgroup analyses which can help identify distinct cognitive mechanisms and variations in parameter ranges across populations. In this study, we focused on model selection to determine which Pavlovian mechanisms best explain behaviours in the Go/NoGo task. The process involved multiple steps to identify candidate models, narrow the model space, and evaluate the winning models.

First, potential mechanisms to model Pavlovian biases in the Go/NoGo paradigm were identified. This step involved reviewing existing literature to select relevant model families and aligning the models with the behavioural hypotheses under examination. Insights from preliminary behavioural data analyses were also considered to ensure the chosen models capture key cognitive processes. (see).

Next, the selected models were individually, and non-hierarchically, fit to the data using minimally informative priors. This step helps verify whether the mechanisms underlying each model are supported by the observed behaviours of individuals in the population. Models that show little evidence of explanatory power were eliminated at this stage, which further justifies the use of an uninformative prior for model identity. Model responsibilities were computed based on the individual fit quality, as captured by the likelihood,

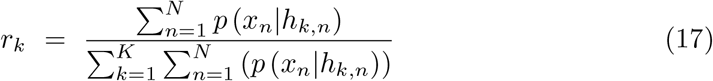

where *h*_*k,n*_ are the fitted parameters for the kth model and nth subject.

No penalty for the number of parameters was applied at this pre-selection step. The goal was to obtain an initial assessment of the plausibility of proposed mechanisms, not to select between models to identify the most parsimonious mechanism capturing the highest amount of variance. Assuming a reasonable selection of models with the similar numbers of parameters, this pre-selection analysis helped to eliminate implausible candidate models.

Parameter recovery was performed for each model in the full model space, based on individually fitted posterior distributions. For this purpose, we generated parameters for one synthetic individual from the posterior distributions of each fitted participant.

Parameter recovery was based on individually fitted models. This naturally accounts for the between-subjects covariance between parameters, ensuring that parameter recovery is tested in relevant parameter ranges. The quality of recovery was assessed in terms of intraclass correlation (ICC).

Following this, the empirical HBI fitting procedure was applied to assess the overall population fit and determine the winning model. The model and exceedance probabilities were calculated to evaluate whether a single model could explain the behaviour of the entire population or if different models would be required for subgroups exhibiting distinct strategies or cognitive processes. Depending on the results and the specific research question, either the eHBI-fitted model would be used directly for subgroup or individual-level analyses, or the entire population would be refitted with the winning model to enable population-level inference and phenotyping.

Model recovery was performed for the selected model subspace. Data was simulated with each model, based on the group posteriors obtained by hierarchically fitting the model to the according dataset. We then ran the eHBI procedure with the full subspace on the simulated data to examine whether the procedure would recover the generative model.

Finally, further subgroup analyses may be conducted to explore emerging patterns within the population. These subgroups may reflect different problem-solving strategies captured by distinct models or variations in cognitive function levels represented by the same model with different parameter values. Such analyses can provide deeper insights into cognitive phenotypes within the studied population.

## Results

We analysed choice data from two distinct populations, who performed different versions of the orthogonalized aversive Go/NoGo task with primary auditory feedback. For each dataset, we first identified candidate models based on previous literature (section 4.3.2). We then refined the model space for the empirical Hierarchical Bayesian Inference (eHBI) procedure by assessing the evidence supporting the prevalence of each mechanism to individuals within the population (section 5.2). The eHBI procedure was subsequently applied to investigate how Pavlovian mechanisms shaped behaviours across groups of participants (section 5.3).

Post-hoc analyses of the winning model(s), alongside correlational analyses between computational mechanisms and behavioural metrics, provided deeper insights into the strengths and limitations of the models within the context of the populations and cognitive task. Additionally, model and parameter recovery analyses, as well as face validity checks, were conducted to ensure the robustness of the findings (see supplementary materials).

### Model-agnostic analyses of datasets I and II

A preliminary model-agnostic analysis was conducted to examine choice behaviour and performance patterns across both datasets (Figure 3). This analysis focused on assessing accuracy for each cue type and evaluating performance trends over trials. For Dataset I, we also investigated the effects of cue-action-outcome reversals on task performance.

**Fig 3.**
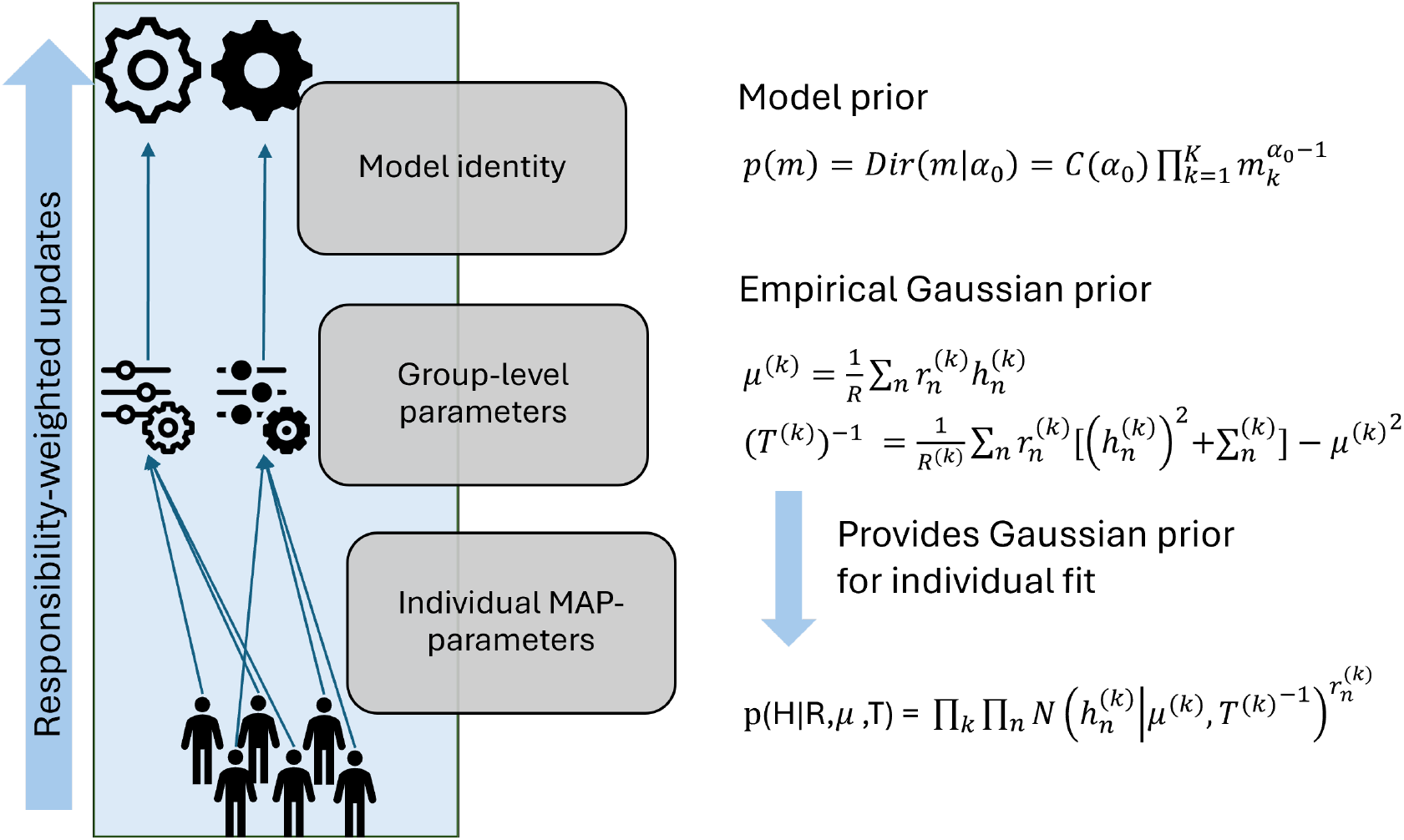
Schematic of the 3 hierarchical levels of the empirical HBI fitting procedure. *At the highest level, the procedure iteratively estimates the model identity for a subject, given current group priors and individual model fits. Model identity is updated as a Dirichlet distribution, which acts as the conjugate prior for the multinomial model, where the model space is given by M* = {*m*_1_, *m*_2_ … *m*_*K*_} . *C*(*α*_0_) *is the normalizing constant for the Dirichlet distribution. At the second level, group priors, with µ*^(*k*)^*as group mean for model k and T* ^(*k*)^ −*as inverse group variance matrix, are updated based on the responsibility-weighted individual maximum a-posteriori (MAP) parameter estimates (with* 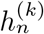 *being the MAP parameters for subject n, fitted by model k, and* 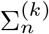 *being the posterior variance of parameters for subject n, fitted by model k, and* 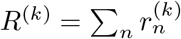*). Model probabilities are given by* 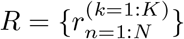 for each model k and individual n, based on relative Bayesian model evidence 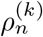. *At the lowest level, individual parameters* 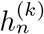 *are estimated, given the updated responsibility-weighted group prior*.

In Dataset I, participants achieved an average accuracy of 68.2% (SD = 10.7%). An ANOVA with accuracy as the dependent variable and condition (Escape vs. Avoid), response type (Go vs. NoGo), and reversal phase (before vs. after) as factors revealed significant main effects of response type and reversal (p=9.7e-15 and p = 2.7e-13, respectively). Accuracy was higher for Go responses and before the reversal phase. Additionally, significant interaction effects were observed between response type and condition (p = .039), suggesting that task performance was modulated by task context. Performance biases in Dataset I displayed high variability. The Go-to-Escape bias (GE) was .206 (SD = .205), while the NoGo-to-Avoid bias (NGA) was -.107 (SD = .221). Although a dominant Go bias was present at the population level across conditions, a significantly lower Go bias was observed in the Avoid condition (p = .004). This indicates the presence of a Pavlovian Inhibitory Avoidance bias in addition to a more pronounced overall active Go-Bias in this dataset.

For Dataset II (Millner et al., 2019), following exclusion criteria, the average accuracy was higher, reaching 76.4% (SD = 15.3%). ANOVA results indicated a significant main effect of response type on accuracy (p = 4.4e-11). The condition-by-response interaction did not reach statistical significance (p =.07). Participants demonstrated better performance for Go-to-Escape cues compared to NoGo-to-Escape cues, while the opposite trend was observed for the Avoid condition. The Go-to-Escape bias of .197 (SD = .257) was similar to Dataset I, while the NoGo-to-Avoid bias of .056 (SD = .266) was more pronounced. As in Dataset I, substantial variability in performance biases was noted across participants.

The observed differences in accuracy between datasets likely reflect differences in task complexity, with Dataset I’s reversal phase contributing to lower overall performance. The high Go-to-Escape bias is consistent across both datasets, suggesting that negatively valenced Go-NoGo tasks with primary feedback capture Pavlovian active-escape response reproducibly. The divergence in NoGo-to-Avoid biases between the datasets highlights a less pronounced and potentially less consistent effect of inhibitory avoidance. While Pavlovian tendencies have previously been associated with suicidality, most literature has focused on active escape or decreased inhibitory biases (Dombrovski & Hallquist, 2021; Interian et al., 2020; Millner et al., 2019; Myers et al., 2023). The discrepancy in findings underscores the need for further investigation into the mechanisms underlying inhibitory control in aversive contexts.

### Model Space Pre-Selection Results

To initialize our model-based analyses, we systematically narrowed the model space to retain only those with significant evidence of capturing relevant mechanisms in the population. This process involved evaluating the average individual fits for each model across both datasets (Figure 5).

**Fig 4.**
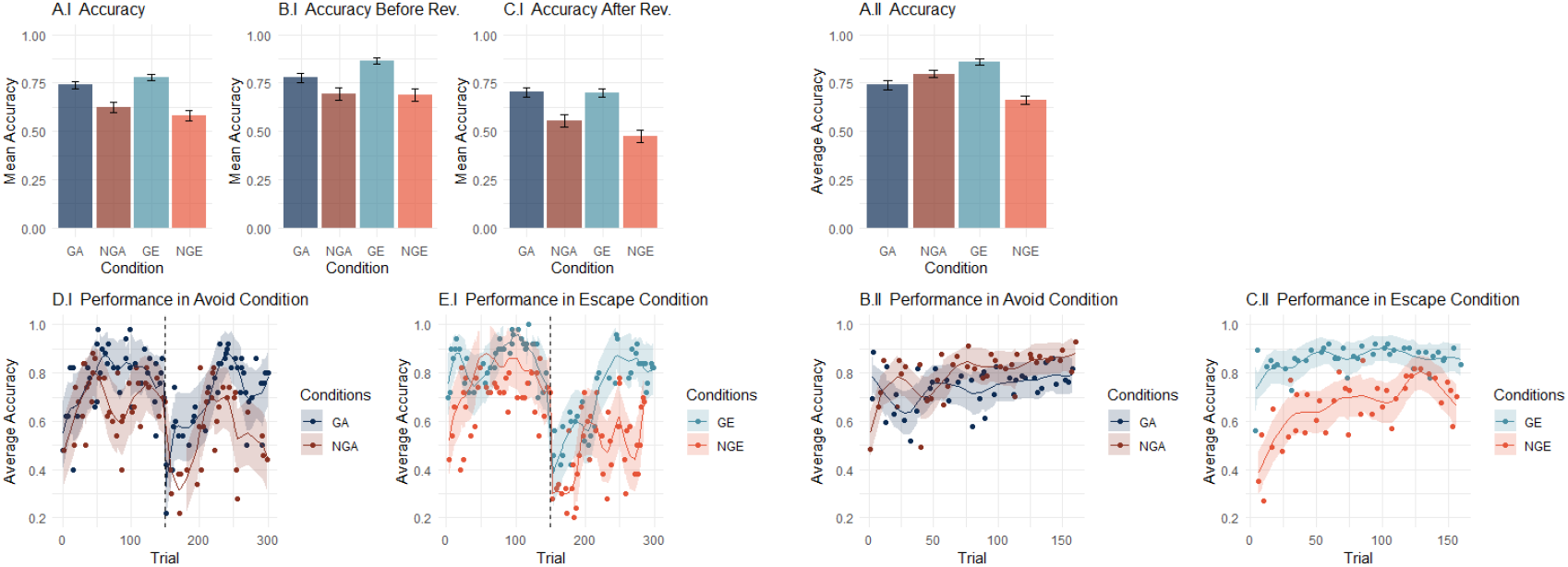
Accuracy of performance on the tasks. The left panels (A.I-E.I) correspond to dataset I. (A.I-C.I) Overall accuracy and accuracy before and after Reversal, split by condition. Task performance is dominated by a Go-bias, where accuracy was higher for Go than NoGo cues during both conditions. (D.I-E.I) average accuracy per trial (dots; the lines show the result of spline smoothing, separately applied before and after the reversal). Accuracy initially increases across conditions, and we observe a clear decline immediately after the cue-outcome reversal, followed by noisy learning. The right panels (A.II-C.II) correspond to dataset II. (A.II) Overall accuracy, split by condition. An effect of response-by-condition can be seen where accuracy is increased for Go-to-escape cues compared to NoGo-to-escape cues, but an inverse effect in the Avoid condition. (B.II-C.II) average accuracy per trial (dots; the lines show the result of spline smoothing). Accuracy increases across conditions over time and plateaus towards the second half of the task. Error bars represent standard error of the mean and shaded areas indicate a 95% confidence interval.

**Fig 5.**
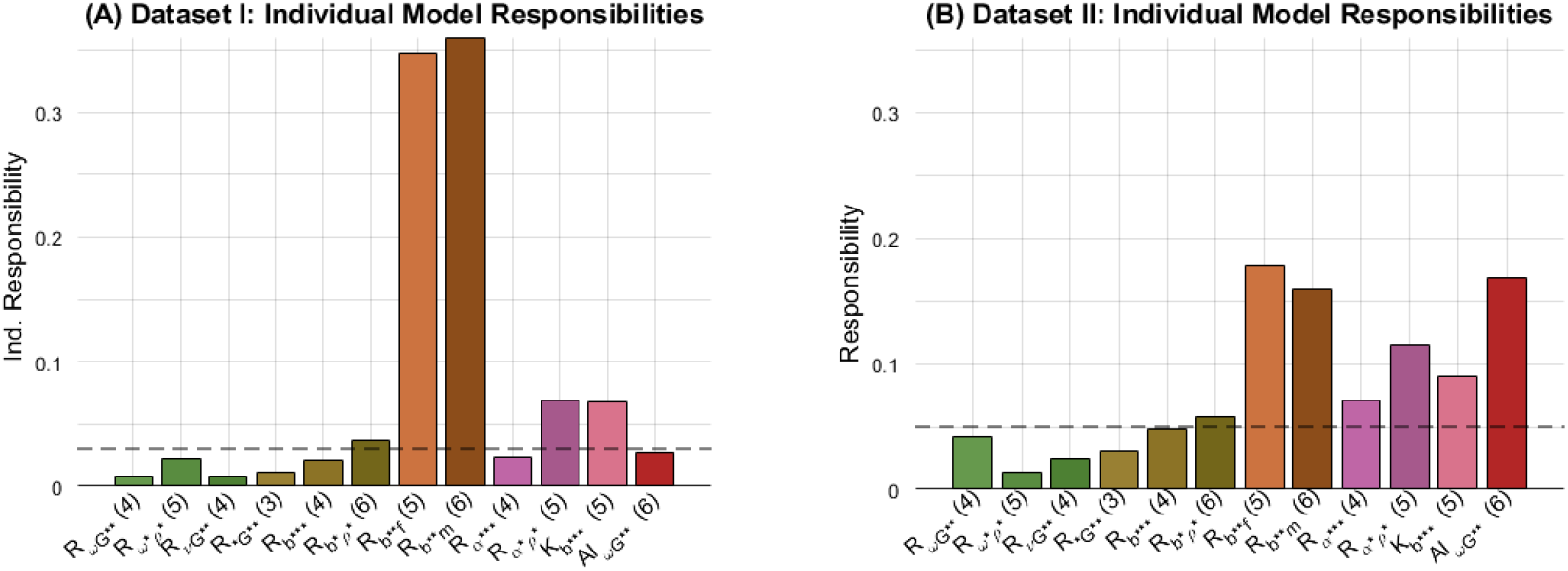
Relative individual model responsibilities. Relative Model responsibilities associated with the full model space individually fitted to each individual in the full population for dataset I (A) and II (B), respectively. A total of 12 different models was fitted to each individual, using non-informative priors. The relative responsibilities were averaged across individuals to obtain a mean relative responsibility of each model for the studied populations. Dashed lines indicate the cut-off for each population to consider a model in the reduced model subspace. Thresholds were chosen empirically and adjusted to create a model space where the least prominent model would have a residual exceedance probability below .01. This approach was chosen to ensure a minimally biased fitting procedure, in which all models with the potential to explain a marked amount of behavioural variance were included.

For Dataset I, two models – *R*_*b*\*\**f*_ and *R*_*b*\*\**m*_ – emerged as dominant, with fit correlations of r = .34 and r = .36, representing the best fit for 18 and 19 participants, respectively (Figure 4). Additionally, the *R*_*α*\**ρ*\*_ model with reward sensitivity and the *K*_*b*\*\*\*_ filter with its uncertainty-adjusted learning rate surpassed the initial threshold of r *>* .05, warranting their inclusion in the final model space. A simple RL model with reward sensitivity was also incorporated after adjusting the threshold to t = .03. The threshold was adjusted after the initial empirical results to create a model space in which the least prevalent model would have a residual exceedance probability below .01, ensuring high confidence in the completeness of the subspace. Models incorporating Pavlovian policies, whether Active Inference, or simpler reinforcement learning (RL) policies, showed minimal evidence of applicability.

In Dataset II, model fits exhibited greater variability, with a broader range of candidate models capable of capturing the behavioural patterns. Consistent with Dataset I, evidence for policy-based RL models was minimal. However, RL models incorporating Pavlovian context biases met the threshold of t = .05 and emerged as potential candidates, except for the simplest versions with only general Go or Pavlovian bias parameters. Due to the more distributed responsibility among models, the default threshold was maintained. In addition, the evidence bias models received significant support. These models feature Pavlovian evidence bias parameters, which are very hard to overcome on the timescale of the task. Additionally, the Active Inference Suicide model (*A*_*ωG*\*\*_) showed strong support at the individual level for this population, representing the only model with Pavlovian tendencies encoded as polices with substantial evidence of relevance.

The differences in model selectivity between datasets may stem from variations in task design. Dataset I, comprising 300 trials, included a cue-outcome reversal, introducing a secondary learning phase that enriched the choice data. This additional complexity likely facilitated model selection by generating distinct learning and relearning windows for each participant. Greater task complexity appears to have enhanced the ability to differentiate between competing models, providing a more comprehensive evaluation of underlying mechanisms.

### Evaluating Model Fit at the Population and Individual Levels

This individual within-participant view of model performance 5 is important, as population-level assessments can hide individual differences. 6 provides insight into models’ ability to predict observed behavioural choices across the population. While relative model responsibility identifies which models best explain individual performance, absolute model fit measures how well a model predicts choice probability across all individuals.

In **Dataset I**, while relative model responsibilities indicated high model selectivity (in favour of *R*_*b*\*\**f*_ and *R*_*b*\*\**m*_), absolute model fit on the population level showed increased fit quality only for *R*_*b*\*\**f*_ , with all other models exhibiting similar absolute fit, but with high within-model variance.

In **Dataset II**, absolute fit probabilities were higher overall but showed greater variability within and across models (Figure 6B). Four models, *R*_*b*\*\**f*_ , *R*_*b*\*\**m*_, *R*_*α*\*\*\*_ and *K*_*b*\*\*\*_ had the highest average absolute fit quality with no clear winner. The large within-model variance of absolute fit (e.g. *A*_*ωG*\*\*_ model in Figure 6B) highlights the importance of assessing model fit at the individual level. Across both datasets, individual level evaluations provided more nuanced insights for model selectivity for individual performance. The support for models above the responsibility threshold in each dataset reflects their superior fit quality compared to other models within individuals.

**Fig 6.**
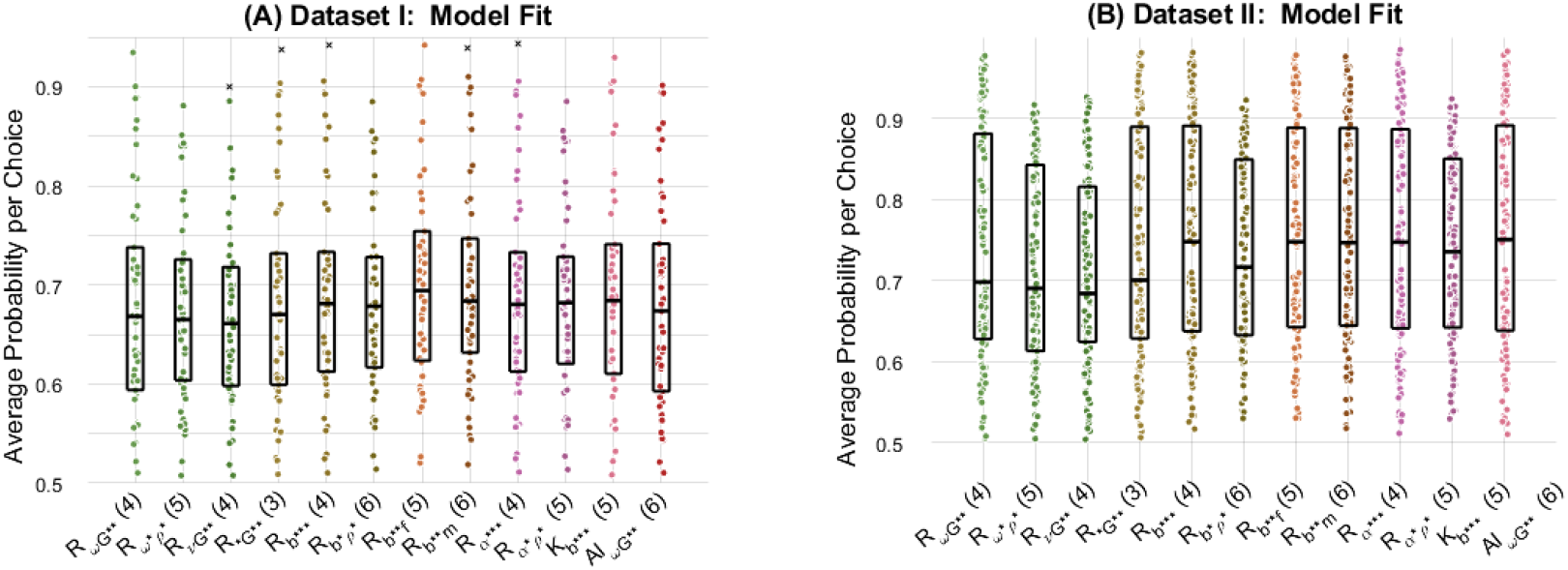
Absolute fit of individually fitted models. Absolute fit of individually fitted models for dataset I (A) and II (B). The average probability per choice for each participant (y-axis) is indicated for each model (x-axis). Boxplots indicate the median, 25^th^ and 75^th^ percentile of the population fit.

Each selected model underwent parameter recovery testing and face validity checks to ensure reliable performance within the relevant parameter ranges based on individual fits (see Section 5.4).

### Replicating Behavioral Patterns with the Best-Fitting Models

Finally, we examined the ability of the best performing models to replicate average choice probability by trial. In Dataset I, RL models with Pavlovian context biases and additional mnemonic mechanisms stood out. The addition of the forgetting factor allowed the model to fit response probability spikes following unexpected feedback more accurately than the *R*_*b*\*\*\*_ model, which lacks mnemonic mechanisms (Figure 7). The working memory model, while capturing broader patterns, lacked this same sensitivity to recent feedback and overshot the average probability in Go-conditions (Figure 7). A similar pattern emerged in Dataset II (Figure 8). Additionally, the Active Inference model garnered significant support at the individual level. Compared to the best-performing RL model with forgetting, the AI model showed slightly reduced accuracy in tracking dips in average population performance throughout the task. The value decay mechanism of the *A*_*ωG*\*\*_ model, based on state-action-prediction-errors, was less sensitive to unexpected feedback compared to the RL model. (Examples of individual trajectories for the best and worst fitting models can be found in Figure S4 and S5)

**Fig 7.**
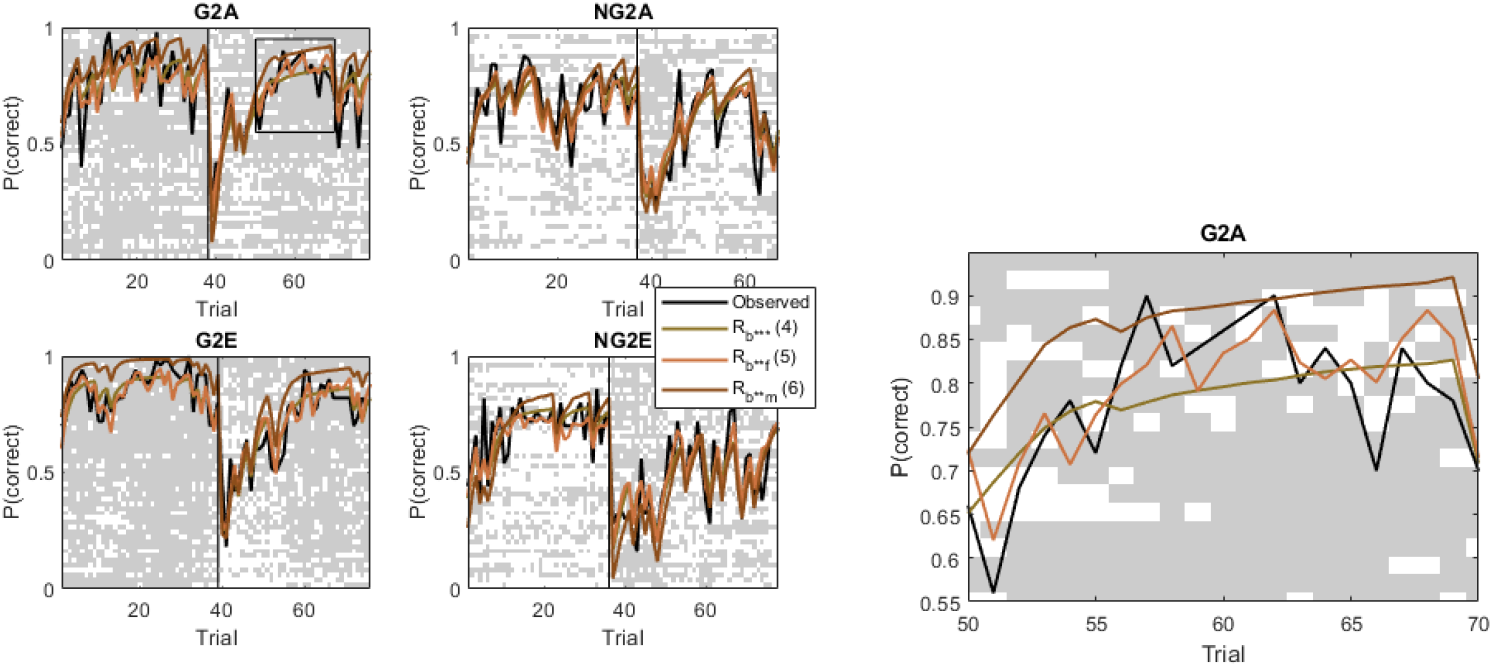
Behavioural fit of best fitting models for dataset I. The plots show the average probability of taking the correct choice for each condition (Go to Escape (G2E), NoGo to Escape (NG2E), Go to Avoid (N2A), NoGo to Avoid (NG2A) across participants in black, compared with the modelled probabilities for the R_b***_ model and its variants with mnemonic mechanisms. The backdrop shows the individual choices of all 50 subjects (rows), sorted by accuracy in the Go-to-Escape condition, where grey indicates a NoGo action and white indicates a Go action. On the right, a zoomed insert is provided.

**Fig 8.**
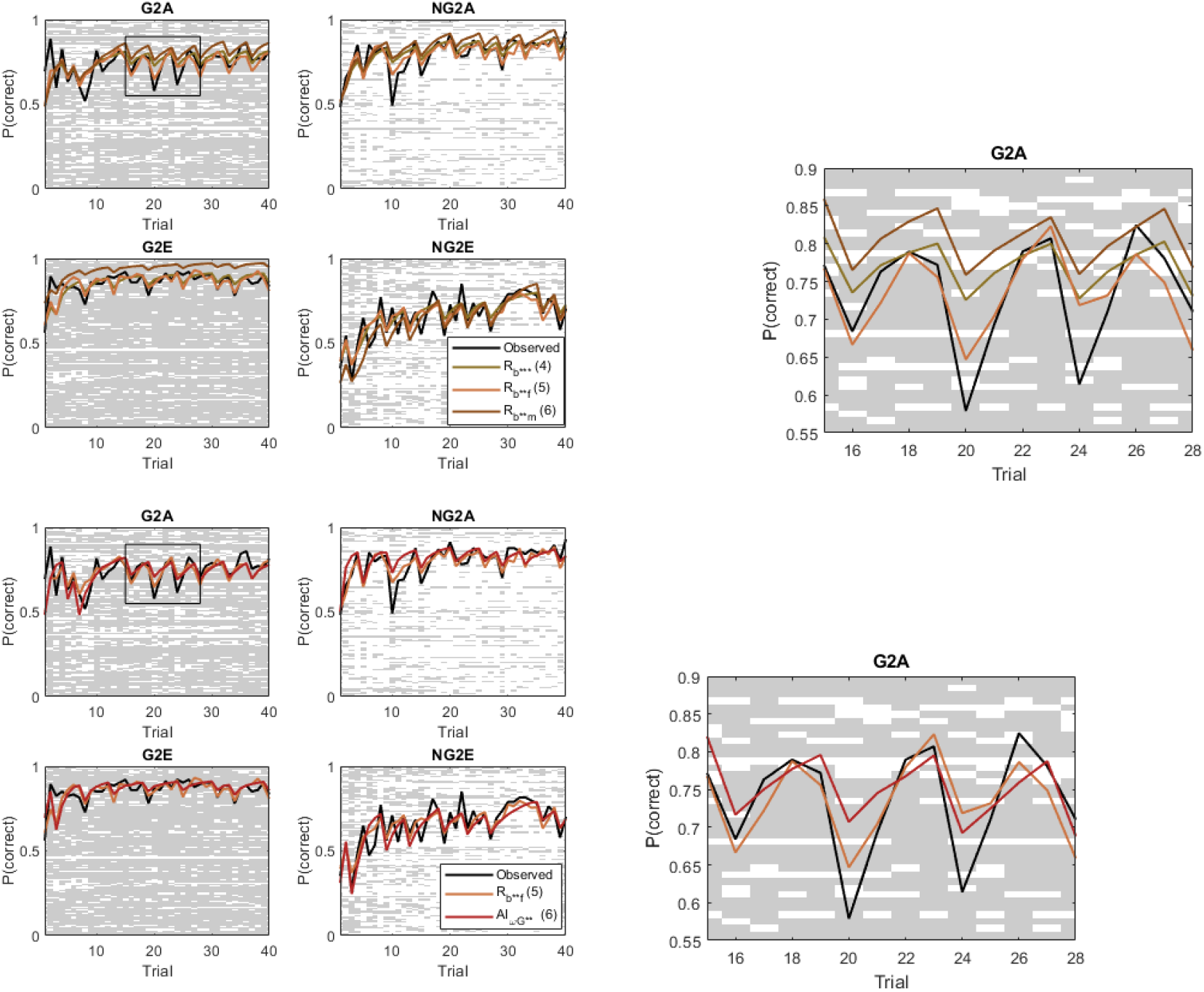
Behavioural fit of best fitting models for dataset II. The plots show the average probability of taking the correct choice for each condition (Go to Escape (G2E), NoGo to Escape (NG2E), Go to Avoid (N2A), NoGo to Avoid (NG2A) across participants in black, compared with the modelled probabilities for the R_b***_ model and its variants with mnemonic mechanisms (top) and the best performing RL model compared to the A_ωG**_ model fit (bottom). The backdrop shows the individual choices of all 114 subjects (rows), sorted by accuracy in the Go-to-Escape condition, where grey indicates a NoGo action and white indicates a Go action. On the right, a zoomed insert is provided.

In simulations, the *A*_*ωG*\*\*_ model clearly outperformed the *R*_*b*\*\**f*_ model in replicating accuracy and Lose-Switch behaviours (*A*_*ωG*\*\*_: Lose-Switch-ICC = .837, Acc-ICC = .973; *R*_*b*\*\**f*_: Lose-Switch-ICC = .7, Acc-ICC = .923), indicating a strong ability to capture some key behavioural traits (see Table S8).

### Model Comparison Results

#### eHBI

The empirical Hierarchical Bayesian Inference (eHBI) procedure was applied to the selected model subspaces for each dataset to analyse responsibility distributions and identify the winning models (Figure 9).

**Fig 9.**
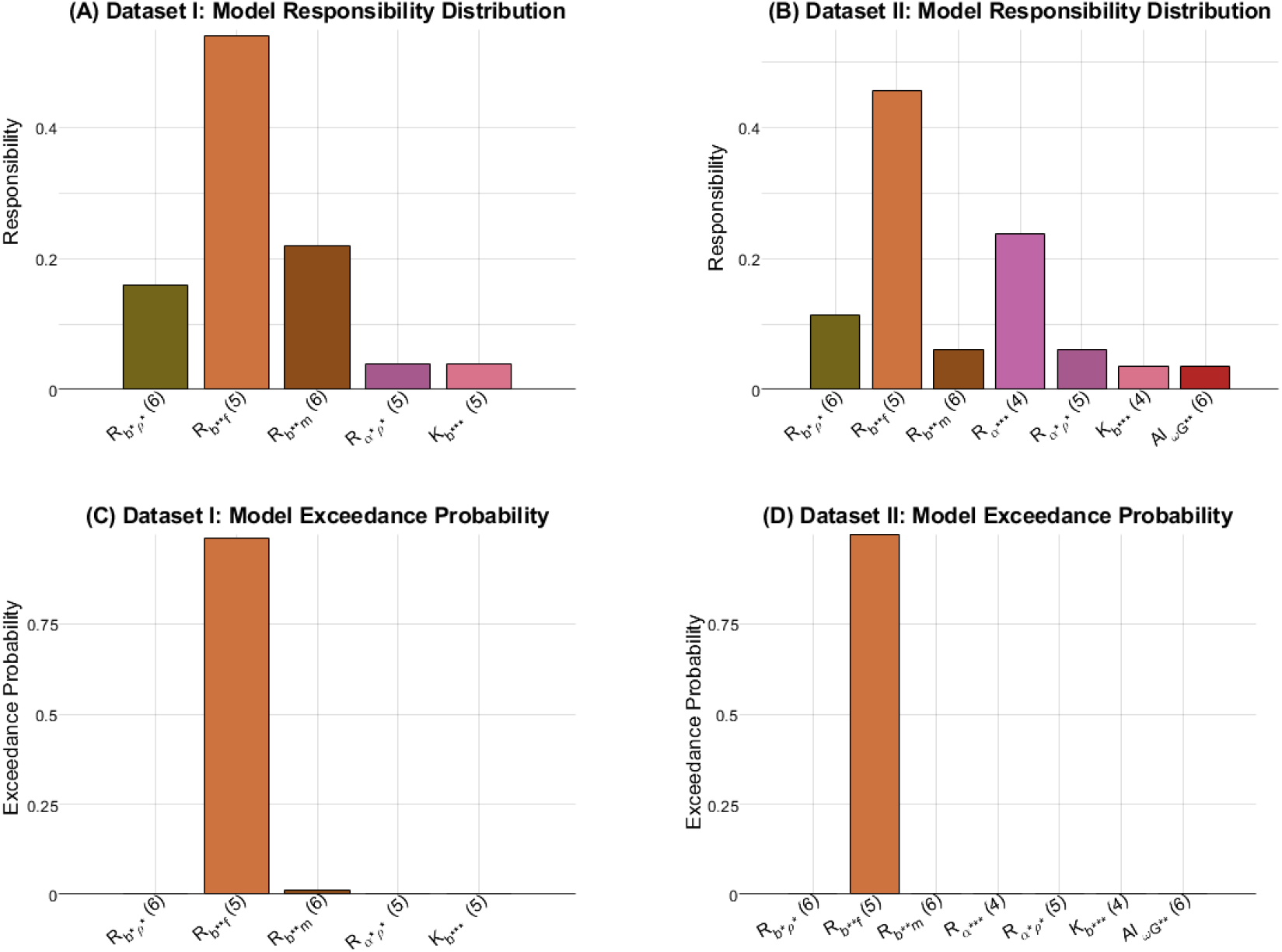
eHBI model comparison results for dataset I and II. *(A) and (B) show the responsibility distribution for the entire population across the respective model subspaces. For dataset I, model responsibilities indicate both the R*_*b*\*\**f*_ *with r =* .*54 and the R*_*b*\*\**m*_ *model with r =* .*22 explain the data very well, followed by the R*_*b*\**ρ*\*_ *model with r =* .*16, For dataset II, again, the R*_*b*\*\**f*_ *model has the highest responsibility with a prevalence of r =* .*46, followed by the simple R*_*α*\*\*\*_ *model with r = 24. The remaining models have low prevalence across subjects. (C) and (D) show the exceedance probabilities for each model for dataset I and II, respectively. Results are consistent across both datasets, indicating the RL model with Pavlovian* context *bias parameters and a forgetting parameter clearly outperforms the competing candidate models on the population level*.

For Dataset I, the *R*_*b*\*\**f*_ model with Pavlovian context biases emerged as the winning model. Consistent with individual-fit evidence, this model accounted for the largest share of model responsibility across participants, with r = .52 and an exceedance probability of *p*_*e*_ *>* .99 at the population level. The *R*_*b*\*\**f*_ model significantly outperformed other candidate models in the subspace. The *R*_*b*\*\**m*_ model was the second most prevalent, capturing less than half the responsibility (r = .24) and best describing only 8 out of 50 participants. Together, these results confirm the importance of accounting for forgetting mechanisms. This opens possibilities to explore the effects of such mechanisms on behaviour further, as previous work has suggested impaired WM capacity in these populations (Richard-Devantoy et al., 2015; Zetsche et al., 2018).

Similarly, the *R*_*b*\*\**f*_ model dominated Dataset II, with an accumulated responsibility of r = .49 and an exceedance probability of *p*_*e*_ = .98. The second highest-scoring model was the simplified *R*_*α*\*\*\*_ model, which accounted for r = .22 of the responsibility.

The responsibility distributions derived from the eHBI model selection deviated notably from the individual pre-selection assessments. Two primary factors contribute to this shift.

First, the eHBI procedure incorporates a parameter penalty term in the model evidence, which was excluded during the initial pre-selection. Models with greater parameter complexity must explain a substantial portion of the remaining data variability to achieve higher responsibility scores. This accounts for the reduced prevalence of the *R*_*b*\*\**m*_ model in the eHBI selection compared to individual-level pre-selection. The *R*_*b*\*\**f*_ model explains similar behavioural effects through a forgetting mechanism [11] but requires one fewer degree of freedom than the *R*_*b*\*\**m*_ model. The design of the task is not optimized to capture working memory processes (as in [11], making the differential treatment of forgetting and working memory unjustified in this context. A similar trend was observed in Dataset II, where the *R*_*α*\*\*\*_ model with reward sensitivity initially scored higher at the individual level but was penalized during eHBI selection due to increased parameter complexity.

Second, the hierarchical fitting in the eHBI procedure evaluates the population, as well as the individual, fit. Fitted group priors impose soft constraints on individual fits, penalizing model fits with parameters deviating from the group prior by reducing their model evidence. If the behavioural patterns of a population are not well captured within a constrained parameter range, a model may exhibit low prevalence at the population level despite demonstrating strong individual fits under uninformative priors. The *A*_*ωG*\*\*_ model for Dataset II exemplifies this phenomenon, as it achieved high individual model evidence but exhibited lower prevalence in hierarchical model selection (see Table S4).

These findings underscore the importance of considering both individual and hierarchical model assessments in behavioural model fitting, as differences in model complexity and parameter variance can significantly impact the selection process and subsequent interpretations.

#### Post-Hoc

To further investigate differences and limitations among the candidate models, we explored how each model captured key behavioral measures, spanning both model-free performance biases and model-based indicators of learning strategy.

In **Dataset I**, we observed a pronounced Go-bias that persisted across Avoid and Escape conditions (Figure 10). Participants displayed relatively high Lose-Switch behaviours (relative to dataset II), suggesting heightened sensitivity to negative feedback. Apart from a notable decrease in accuracy, the raw behavioral data did not reveal significant immediate effects of cue-outcome reversal, indicating a general difficulty in re-learning cue-action associations during the second half of the task.

**Fig 10.**
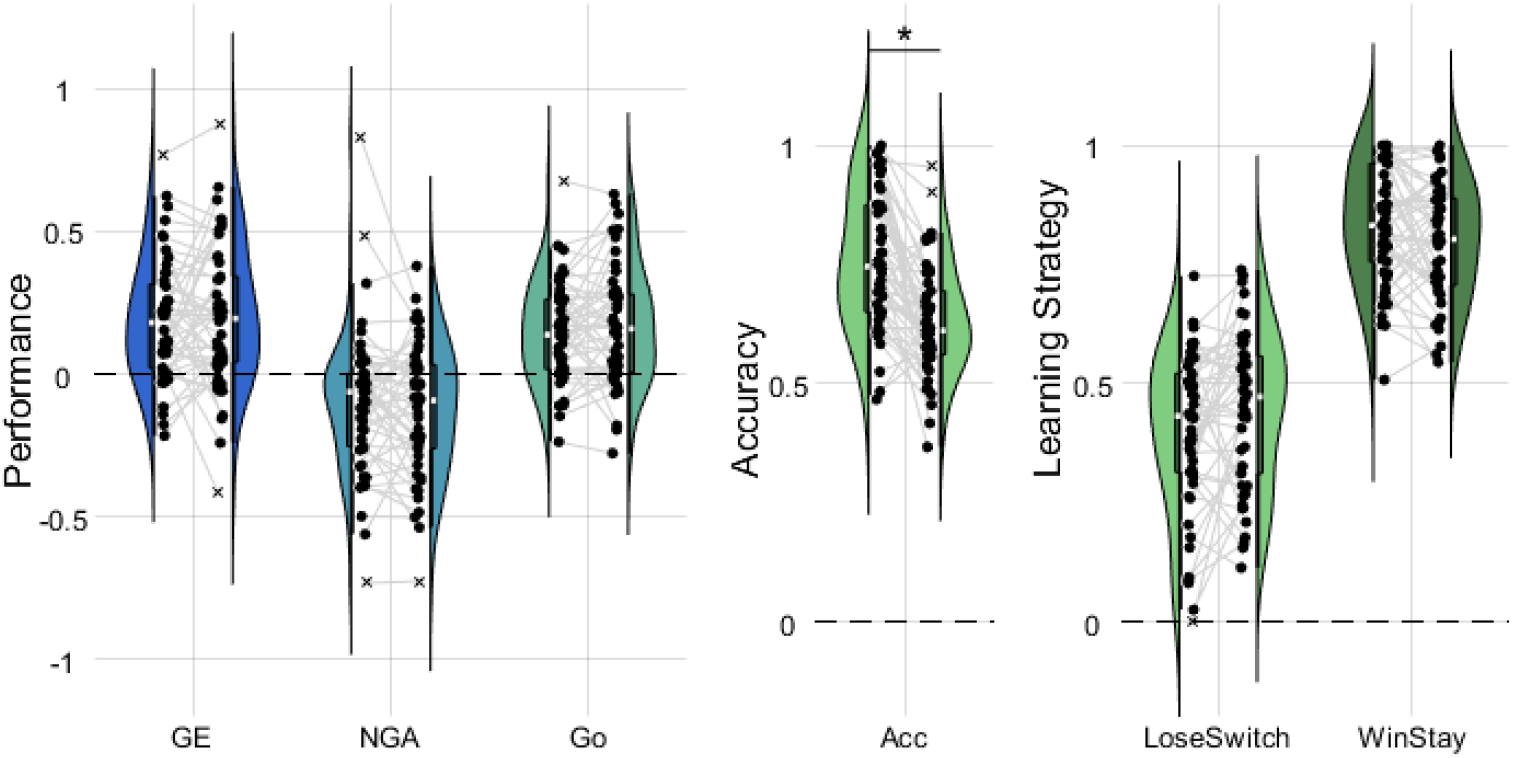
Model-agnostic behavioural measures for dataset I. Model-agnostic behavioural measures for dataset I, including performance biases (Go-to-Escape Bias GE, NoGo-to-Avoid Bias NGA and overall Go-Bias), accuracy and learning strategies (Lose-Switch and Win-Stay behaviours). Each measure is shown before (left half-violin) and after (right half-violin) reversal. The bias in the avoid condition is anti-Pavlovian, meaning participants display a weak Go bias during Avoid conditions. The only measure with a significant difference before and after reversal is accuracy, with a significant decrease in the second half of the task.

Next, we assessed model differences by grouping participants according to their winning model (*max*_*k*_(*p*_*k*_(*m*))) within the eHBI procedure (Figure 11, p-values can be found in Table S5). Models that best fit fewer than four individuals, such as the *K*_*b*\*\*\*_ filter and *R*_*α*\*\*\*_ with reward sensitivity, were excluded from subsequent subgroup analyses to ensure sufficient inferential power.

**Fig 11.**
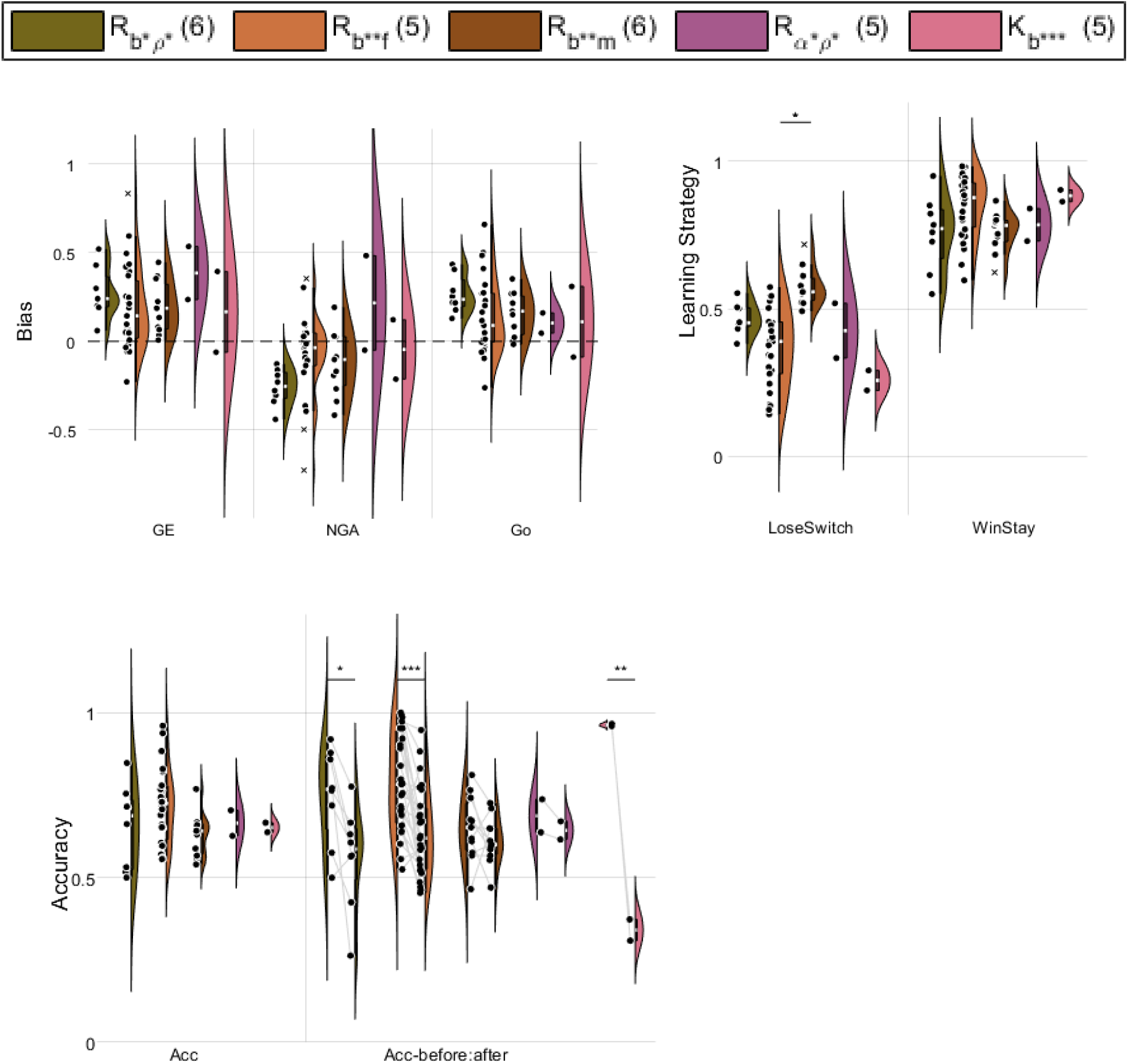
Subgroup model-agnostic behavioural measures for Dataset I. *Subgroup model-agnostic behavioural measures for Dataset I, including performance biases (overall Go-Bias, Excitatory-Escape Bias and Inhibitory-Avoid Bias), learning strategies (Lose-Switch and Win-Stay behaviours) and accuracy (total, before and after reversal). Significant differences between subgroups are indicated by star symbols (FDR-equivalent of \**: FDR_cuttoff = 4.4e-4 *- p<*.*05, **:* FDR_cuttoff = 0.0*- p<*.*01, ***:* FDR_cuttoff = 0.0*-p<*.*001). Significance is calculated according to the FDR-corrected p-value of a two-sided t-test. Significant differences between accuracy before and after reversal are uncorrected, as they are evaluated within each subgroup only*. (see Figure S7 for accuracy measures by condition and Table S5 for p-values)

Within the *R*_*b*\*\**f*_ group, there was substantial variance across most model-agnostic behavioural measures. Although this group showed a decrease in inhibitory avoid bias relative to the *R*_*b*\**ρ*\*_ model (p = .049,) and a decrease in Lose-Switch strategy compared to the *R*_*b*\**ρ*\*_ model (p = .048, *FDR*_*cuttoff*_ = 4.4e-4), as well as an increase in the Win-Stay strategy, these effects did not survive correction. In contrast, the *R*_*b*\*\**m*_ group exhibited significantly higher Lose-Switch behaviour compared to the *R*_*b*\*\**f*_ group (p = 1.6e-5) and showed a trend toward significance relative to the *R*_*b*\**ρ*\*_ group (p = .001).

Despite the limitations imposed by very small sample sizes, certain behavioral traits nonetheless appeared to be better explained by specific models. The *R*_*b*\*\**f*_ model, for example, captured individuals across a wide range of accuracies and revealed a significant effect of the reversal on performance accuracy (Fig.10). This group showed adaptive learning strategies aligned with the probabilistic feedback structure, accounting for some of the highest performers in the dataset. The forgetting mechanism of the *R*_*b*\*\**f*_ model may help explain task performance by gradually reducing the influence of older information on action selection over time. This hypothesis is supported by the model’s ability to replicate performance accuracy, particularly before the reversal, as well as Win-Stay patterns (see S, Tab.S10). Given the task design, which involved up to 20 trials between cue repetitions, these findings suggest that explicitly modeling value decay (as in *R*_*b*\*\**f*_) is crucial for accurately capturing performance during long inter-trial intervals.

The *R*_*b*\*\**m*_ model, on the other hand, predominantly captured participants who exhibited increased Lose-Switch behaviour. This model tracks action values using both working memory and instrumental learning systems, placing strong emphasis on the most recent trial outcome. As a result, it demonstrates a high capacity to reproduce Lose-Switch and Win-Stay patterns (Table S8). However, the reliance on perfect recall of recent feedback also introduces volatility in action-value estimates, potentially contributing to lower overall accuracy in a probabilistic task such as Go/NoGo. A significant negative correlation between the working memory contribution (*ρ*_*W M*_) and accuracy following unexpected feedback (R = -.619, p = .042) supports this interpretation.

**Dataset II** showed large variability in performance biases, with minimal Go-bias and Pavlovian performance biases in Avoid and Escape conditions (Figure 12). Overall, participants displayed low Lose-Switch and high Win-Stay behavior, consistent with relatively strong task performance. As in Dataset I, any subgroups with fewer than four individuals were excluded from the analyses.

**Fig 12.**
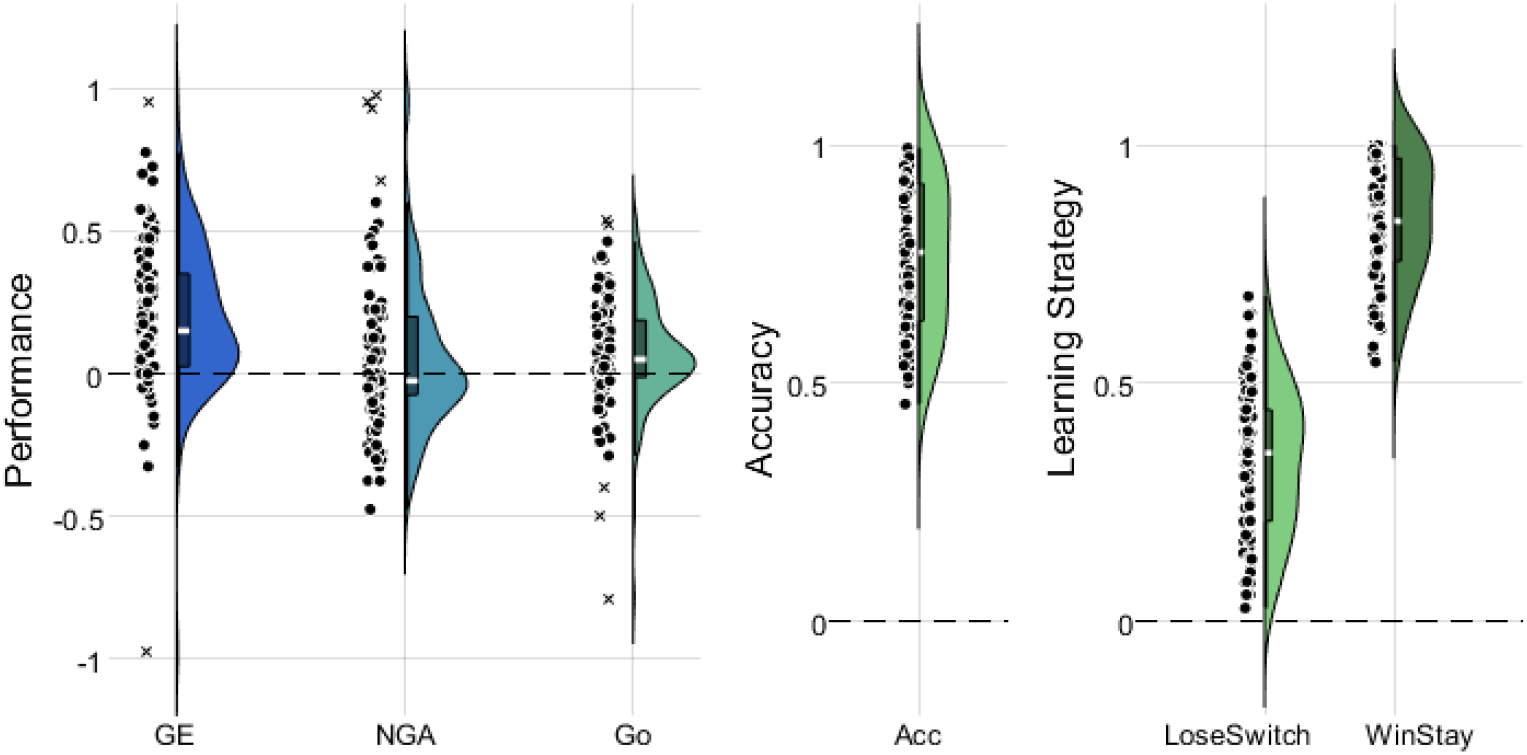
Model-agnostic behavioural measures for dataset II. Model-agnostic behavioural measures for dataset II, including performance biases (Go-to-Escape Bias GE, NoGo-to-Avoid Bias NGA and overall Go-Bias), accuracy and learning strategies (Lose-Switch and Win-Stay behaviours).

Compared to the *R*_*b*\*\**f*_ , *R*_*b*\*\**m*_, and *R*_*α*\*\*\*_ models, the *R*_*b*\**ρ*\*_ model captured a subgroup characterised by significantly lower Go-bias, increased Active Escape, and enhanced Inhibitory Avoid Bias (Figure 13, p-values can be found in Table S6). Differences in learning strategies and accuracy across subgroups were less consistent.

**Fig 13.**
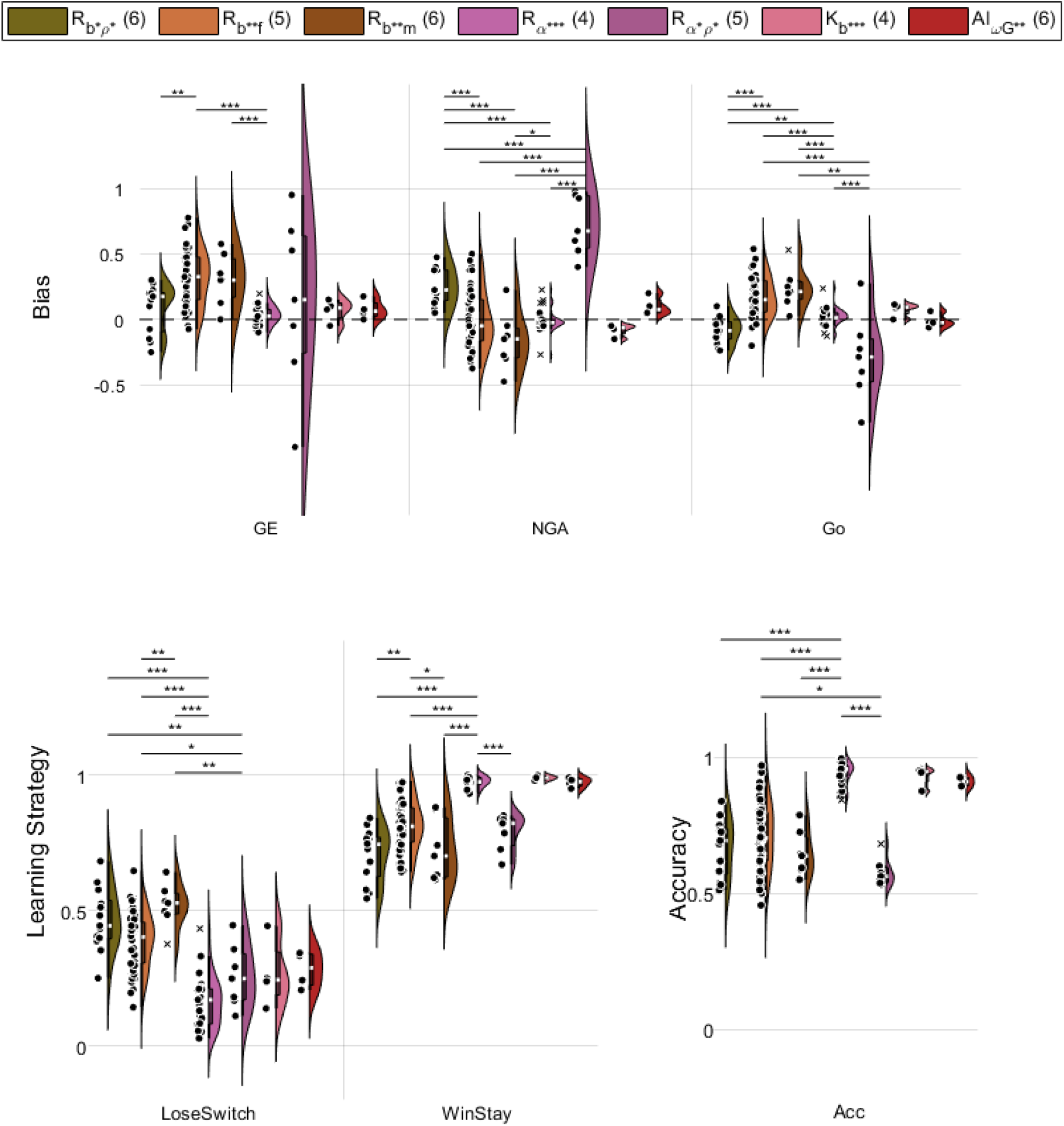
Subgroup model-agnostic behavioural measures for Dataset II. *Subgroup Model-agnostic behavioural measures for Dataset II, including performance biases (overall Go-Bias, Excitatory-Escape Bias and Inhibitory-Avoid Bias), learning strategies (Lose-Switch and Win-Stay behaviours) and accuracy. Significant differences between subgroups are indicated by star symbols (FDR-equivalent of *:* FDR_cuttoff = .021 *- p<*.*05, **:* FDR_cuttoff = .003*- p<*.*01, ***:* FDR_cuttoff = .0002*-p<*.*001). Significance is calculated according to the FDR-corrected p-value of a two-sided t-test*.(see Figure S8 for accuracy measures by condition and Table S7 for p-values)

The *R*_*b*\*\**f*_ and *R*_*b*\*\**m*_ models captured overlapping behavioural profiles, with the primary point of differentiation being the increased Lose-Switch behaviour in the *R*_*b*\*\**m*_ group (p = 0.003). This finding parallels results from Dataset I, where the *R*_*b*\*\**m*_ model excelled at explaining heightened Lose-Switch behaviour, a pattern attributed to the working memory system’s perfect recall of the most recent trial.

The *R*_*α*\*\*\*_ model, representing the third-largest subgroup, was marked by low variance in model-agnostic behavioral measures. Individuals in this subgroup showed minimal performance biases, significantly lower Lose-Switch and Win-Stay behaviour than the other large-model subgroups (*R*_*b*\**ρ*\*_, *R*_*b*\*\**f*_ and *R*_*b*\*\**m*_), and higher overall accuracy. This subgroup thus represents high-performing individuals with few biases, whose behavior is sufficiently captured by a reduced-parameter model. By contrast, lower-performing participants were better described by models incorporating Pavlovian context biases or additional flexibility, such as forgetting mechanisms or reward sensitivity adjustments.

Overall, these analyses suggest that different computational models capture distinct behavioural profiles within the population, shedding light on potential mechanisms underlying performance variability in aversive Go/NoGo tasks. Notably, in both datasets, models featuring forgetting or working memory mechanisms best explained subgroups with stronger Lose-Switch behaviors. Their ability to replicate these learning strategies (Table S8), illustrates how the models’ parametrization, accounting for recency effects, improves fit for individuals whose behavior is particularly influenced by recent outcomes.

### Model Recovery and Face Validity

The winning models for Datasets I and II reproduced behavioral variables with high fidelity, as indicated by intraclass correlation coefficient (ICC) values above 0.7 (Figure 14; Table S8). In contrast, non-winning models showed greater variability, with ICC values dipping as low as 0.29 for the Lose-Switch strategy. Because the Lose-Switch pattern was not explicitly targeted by the model fitting, its lower reproducibility was unsurprising. Overall, these findings point to moderate parameter recovery across models but highlight notable differences in how effectively they capture and reproduce key behavioral features. To ensure the interpretability of these analyses, we conducted model recovery and parameter recovery assessments, as well as face validity checks.

**Fig 14.**
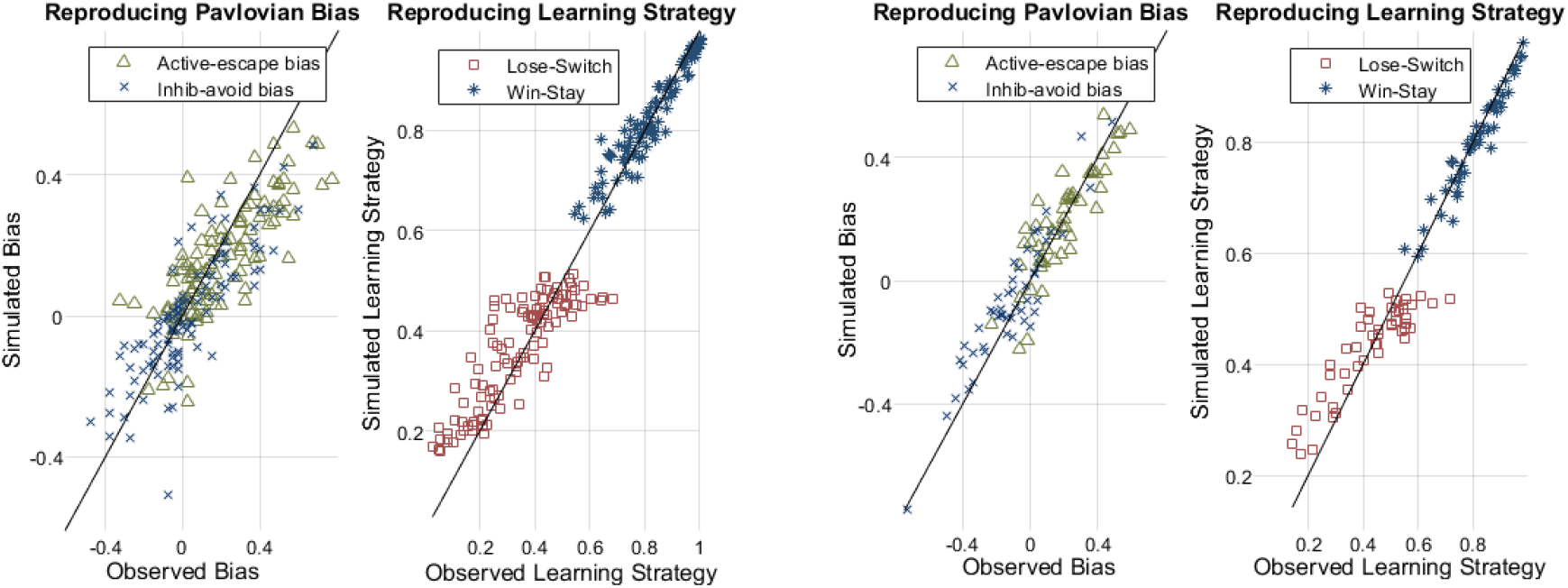
Face Validity of winning model for Datasets I and II. Face Validity of winning model R_b**f_ for Datasets I and II (panel A and B, respectively). For each dataset, the behavioural measures for active-escape and inhibitory-avoid biases and lose-switch and win-stay metrics computed on generated behaviour is plotted against the computed behavioural metrics on the observed data.

A model recovery analysis was performed for the subspaces for both datasets (Figure S8). For Dataset I, the true generative model overwhelmingly dominated in every instance with a median probability of 100% and the lowest probability being 99.3%. The same was true for Dataset II, in which the true generative model dominated with a median probability of 88.6%, except for the *K*_*b*\*\*\*_ model. Data generated by the *K*_*b*\*\*\*_ model was best fitted by the *R*_*α*\*\*\*_ model with one fewer parameter, indicating that the complexity of the Kalman filter was not capturing sufficient variability of the data to justify the additional degree of freedom. This is in line with the results of the eHBI procedure for Dataset II, in which the *R*_*α*\*\*\*_ had significantly stronger support.

Next, we performed parameter recovery for each model in the full model space. Except for the uncertainty parameter *σ* in the Kalman filter, all parameters had ICC values *>*.4, indicating moderate to good recovery (Table S7). The median ICC across all models was *ICC*_*m*_ = .89 (median absolute deviation or MAD = .05) for Dataset I and *ICC*_*m*_ = .84 (MAD = .09) for Dataset II.

To assess the face validity of each individual model, we generated synthetic data for each individual based on parameters from the hierarchically fitted model, and comparing the replication of various behavioral measures -Pavlovian biases (Active Escape, Inhibitory Avoid), learning strategies (Win-Stay, Lose-Switch), and choice accuracy. In both datasets, models in the selected subspaces generally reproduced these measures well, with the notable exception of Lose-Switch behaviour in models that included context-dependent reward sensitivity (*ρ*), which exhibited poor reproducibility. This result underscores the need for caution when using these particular models to study behaviors in greater detail. In contrast, models excluded by eHBI showed mixed face validity, suggesting limitations in their ability to capture the observed behaviors in these populations. (Table S8)

Overall, the parameter and model recovery analyses performed well across tested models, but the face validity results highlight that certain model-agnostic behavioral measures—particularly Lose-Switch—are not adequately captured by many models. These findings underscore the need for careful and nuanced interpretation, especially when drawing conclusions about specific learning strategies from model-based analyses.

## Discussion

In this study, we investigated the cognitive mechanisms underlying performance in negatively valenced Go/NoGo tasks among individuals with suicidal thoughts and behaviors (STB). By adapting and applying the hierarchical Bayesian inference (HBI) procedure [25], we aimed to identify which models best captured Pavlovian and instrumental learning processes in two separate datasets featuring aversive primary feedback. Throughout our analyses, we focused on potential Pavlovian biases and the role of learning mechanisms such as value decay (forgetting) and policy-based adaptations.

### Key Findings

**Dataset I** was characterized by a persistent Go-bias throughout the task, even though we hypothesized that the heightened environmental volatility and potential reversal-learning deficits in suicidal populations [22] would elevate Pavlovian biases following a contingency reversal on trial 151. Our initial hypothesis was that contingency reversal would primarily impact instrumental learning, as instrumental associations are action-dependent and require updating based on new evidence. Conversely, Pavlovian associations, being action-independent, should remain unaffected by the reversal. However, this was not observed. Instead, the reversal procedure (affecting both cue-action associations and auditory feedback) simultaneously challenged both learning systems. Prior research suggests that reversal learning deficits may not be specific to a single learning system [49]. If both systems were similarly impaired following the reversal, the Pavlovian system could not compensate for deficits in instrumental learning, explaining the absence of increased Pavlovian biases post-reversal.

Model comparison using the eHBI procedure indicated a clear prevalence of the *R*_*b*\*\**f*_ model, which incorporates static Pavlovian bias parameters with a decay parameter for learned values, capturing forgetting over time. No evidence of policy-based Pavlovian mechanisms was found. These results suggest that Pavlovian biases do not require significant associative learning before being evident in behavioural responses. The feedback structure, emphasizing primary aversive stimuli, may inherently activate Pavlovian associations, reducing the need for learned associations. Additionally, in this version of the Go/NoGo task, the presence of the aversive sound indicates the context (Avoid or Escape) to the participant right away, minimizing the requirements for associative learning [7, 36]. The additional complexity introduced by a value decay parameter was justified by their superior explanatory power relative to simpler, bias-only models, particularly in the task context involving prolonged inter-trial intervals and probabilistic feedback.

In contrast, **Dataset II** showed stronger Active-Escape tendencies and weaker Inhibitory-Avoid biases, diverging from the dominant Go-bias observed in Dataset I. Similar trends were observed, with *R*_*b*\*\**f*_ emerging as the winning model and minimal evidence supporting policy-based models. Unlike Dataset I, the *A*_*ωG*\*\*_ model gained substantial support during individual model comparison. However, complexity penalties within the hierarchical Bayesian framework limited this model’s overall support, emphasizing the necessity of balancing complexity with explanatory precision.

An important methodological distinction between the two datasets is the number of trials and the presence of a reversal in Dataset I. Dataset I, with 300 trials and a critical reversal point, provided a rich source of behavioral variability, resulting in more pronounced model selectivity. By contrast, Dataset II included only 160 trials, limiting the ability to distinguish among competing models.

The results of this study align with, and extend, prior research investigating Pavlovian tendencies in Go/NoGo tasks. Early work by Guitart-Masip et. al [9] proposed modeling Pavlovian tendencies in the now common reward/punishment version of the Go/NoGo task with secondary feedback as learnable policies but did not explore alternative mechanisms, such as static biases. Adams et. al compared policy- and bias-based RL models, including versions with forgetting and reward sensitivity, as well as an Active Inference model, in a traditional secondary-feedback Go/NoGo task in healthy controls [15]. They found that a context bias RL model with reward sensitivity best explained behavior of most of the population, while adding a forgetting parameter did not improve fit. Importantly, in their study, the Pavlovian bias parameters were assumed to be of equal size across reward and punishment conditions and forgetting was only tested for the best-performing RL model (with reward sensitivity) and did not improve the fit further. Notably the only Pavlovian context bias model they tested performed best in model comparison, resonating with the robust evidence of context biases we identified under aversive primary feedback.

Additionally, research by Swart et al. [36] emphasized the influence of stable biases when the cue indicates the task condition, and [7, 10] demonstrated how aversive primary feedback can elicit immediate Pavlovian reactions. Our results align with these studies: static Pavlovian biases seem to manifest quickly once the aversive context is evident, without requiring extensive associative learning.

### Applications of the eHBI Procedure

The eHBI procedure is particularly valuable for model selection in datasets with heterogeneous population characteristics and unclear cognitive parametrization. Each model and its mechanisms bring specific assumptions and implications about Pavlovian and instrumental learning systems and how these mechanisms influence observed behaviors. By treating model identity as a random factor during fitting, the eHBI procedure facilitates unbiased comparisons and supports subgroup analyses. Such analyses are invaluable when different segments of a population rely on distinct mechanisms, as different models can be used to parametrize each segment accurately. While we did not observe significant model heterogeneity in the studied populations, previous work on larger datasets with different model prevalences has shown the importance of this [15, 43].

Alternatively, when a population relies on the same cognitive strategy but exhibits varying functionality levels, subgrouping analyses can cluster individuals into distinct parameter ranges. This data-driven approach holds promise for sample stratification and diagnostic purposes. For example, clustering may reveal phenotypic subgroups, enabling targeted interventions or more nuanced clinical assessments [43].

## Limitations and Future Directions

While our hypothesis-driven approach clarified the cognitive processes underpinning performance in these suicidal populations, it was inherently limited by the set of proposed models. To uncover more nuanced or previously unrecognized mechanisms, exploratory computational methods, such as the Disentangled Recurrent Neural Networks (DisRNNs; [50]; see also [51]), which autonomously discover interpretable cognitive structures directly from behavioral data, can be employed. Unlike our constrained method, DisRNN employs neural network architectures with disentanglement and information bottlenecks, allowing for the discovery of latent cognitive processes beyond initial assumptions. Integrating such exploratory methods alongside structured hypothesis-driven modelling could uncover subtle or novel cognitive phenotypes, enhancing clinical subtyping and informing targeted interventions in suicidal populations.

Several further limitations must be acknowledged. The relatively small sample sizes, particularly in Dataset I, and differences in task design (e.g., auditory feedback, task length, presence of reversal) limit the generalizability of the findings. Additionally, heterogeneity in demographic factors, psychiatric diagnoses, and medication use further complicate direct comparisons between datasets and are limited to populations with symptoms of suicidality. These factors interact in complex ways to shape Pavlovian and instrumental learning, contributing to variability in responses to reinforcement and aversive stimuli. Psychiatric conditions such as anxiety disorders and PTSD, for instance, have been linked to heightened sensitivity to aversive stimuli, which can impede the extinction of conditioned responses [52]. Additionally, the co-occurrence of depression and PTSD may further dis-regulate learning mechanisms, perpetuating maladaptive behavioral patterns [53]. The current work focused purely on identifying cognitive mechanisms based on behavioural data, meaning the exploration of clinical validity and utility were beyond scope. Future work will address this gap, investigating the connection between computational parameters and clinical features within tested populations. Similarly, developmental and age-related factors also play a crucial role in shaping learning behavior. While younger individuals tend to exhibit greater adaptability in learning new associations, older adults may experience difficulties in unlearning previously learned behaviors due to cognitive decline [54].

Building on these complexities, stratifying participants by demographic, psychological, and medical characteristics can bolster model interpretability and enhance the precision of computational analyses. Future studies should address these limitations by systematically testing task variations in controlled, diverse populations to establish correlations between task-induced behaviors and well-constrained model parameters. Such work could improve task design, refine model selection and optimize parameter recovery.

Finally, this study focused exclusively on binary choice data, excluding reaction time, a continuous variable that could offer deeper cognitive and behavioral insights.

Incorporating reaction time data in future model analyses may enhance understanding of underlying decision-making processes.

Overall, our study underscores the utility of the eHBI procedure for uncovering the cognitive mechanisms underlying behavioural variability in psychopathologies. Future work integrating more nuanced task designs and multimodal data could further advance the understanding of Pavlovian biases and their role in mental health disorders.

In conclusion, our study supports the centrality of stable Pavlovian biases and forgetting mechanisms in decision-making in suicidal populations. Further integration of exploratory computational methodologies, such as DisRNN, alongside structured approaches and subgrouping analyses could deepen cognitive insights.

## Supplementary Information

**Figure S1.**
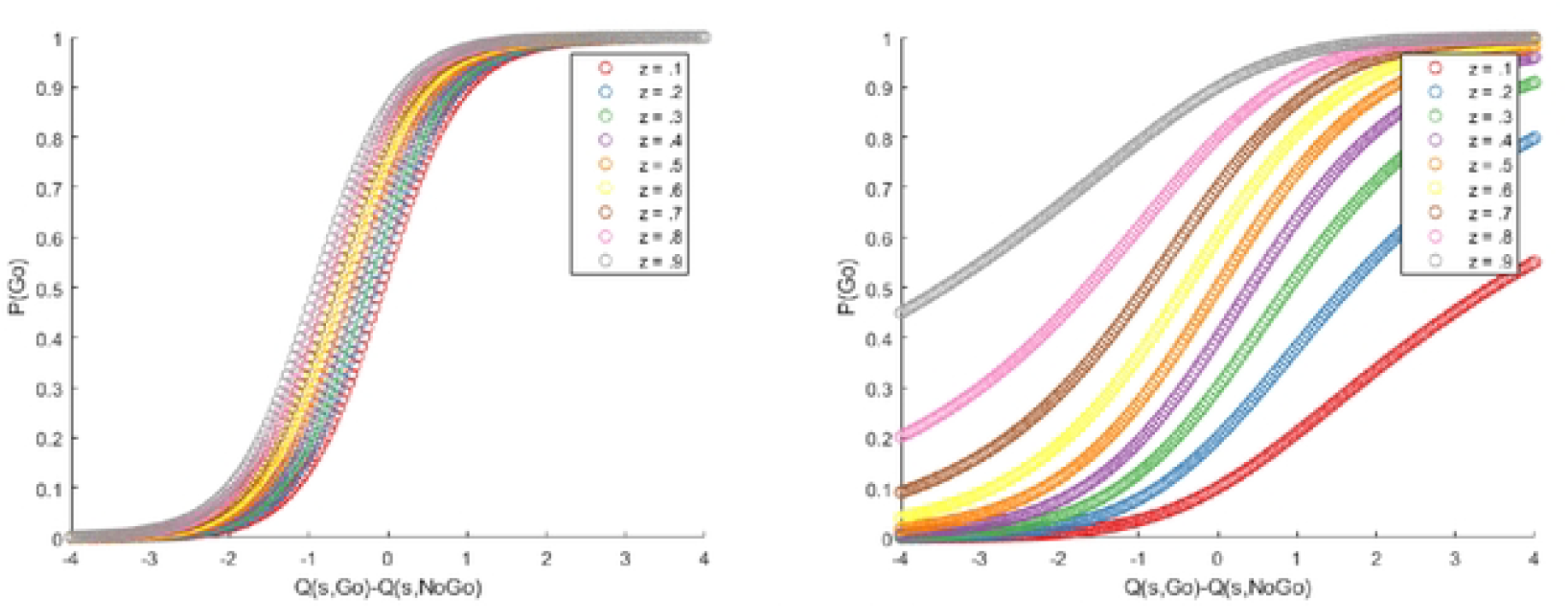
SoftMax function of additive and multiplicative Go biases. Illustrative translation of instrumental values and Pavlovian tendencies into actions via SoftMax functions, as employed in the considered models. On the left, the impact of Pavlovian bias parameters as employed in the R_*α*\*\*\*_ model can be seen. On the right, the impact of static additive Pavlovian bias parameters is shown.

**Figure S2.**
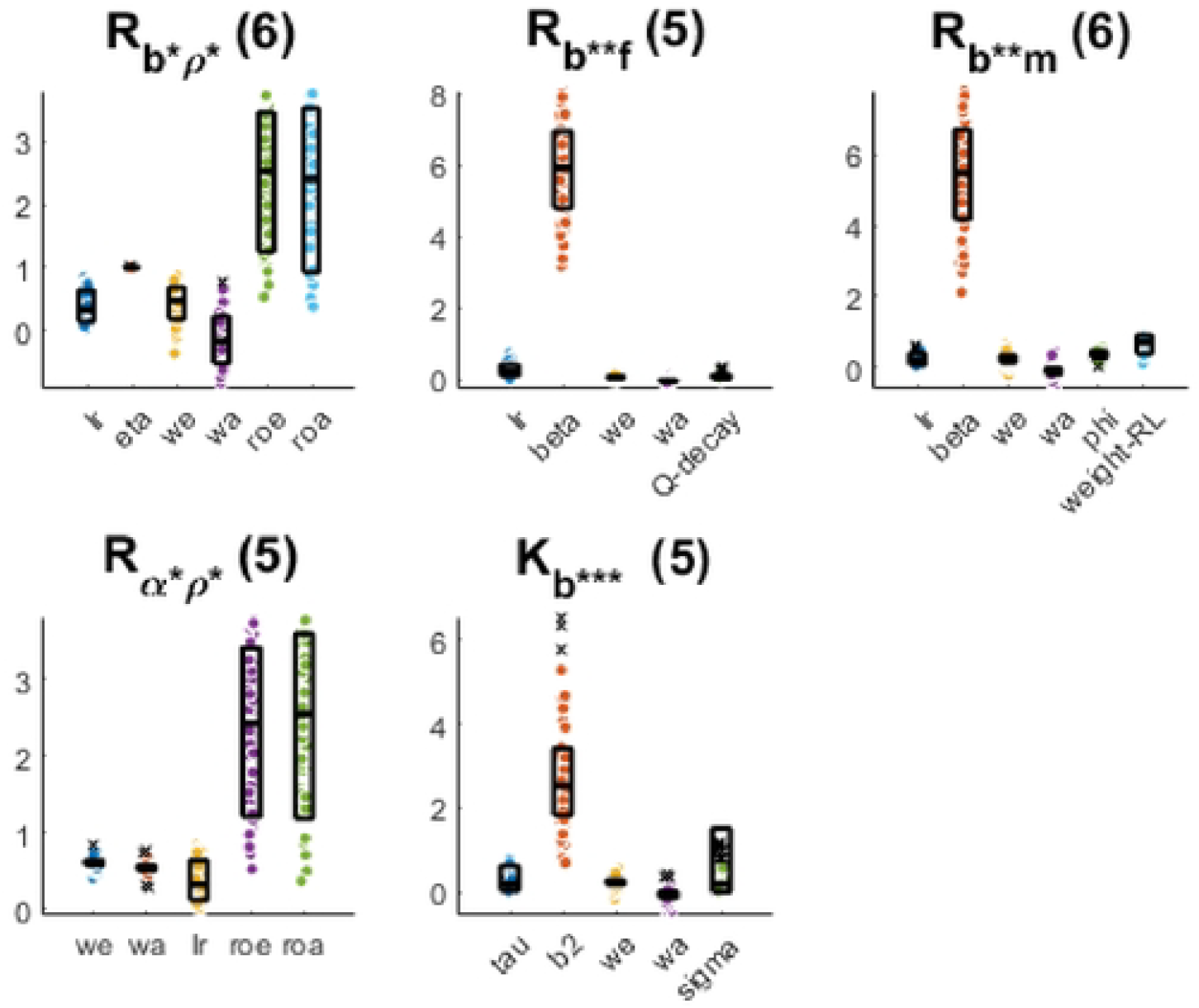
Fitted parameter ranges for individually hierarchically fitted models of the selected subspace for Dataset I.

**Figure S3.**
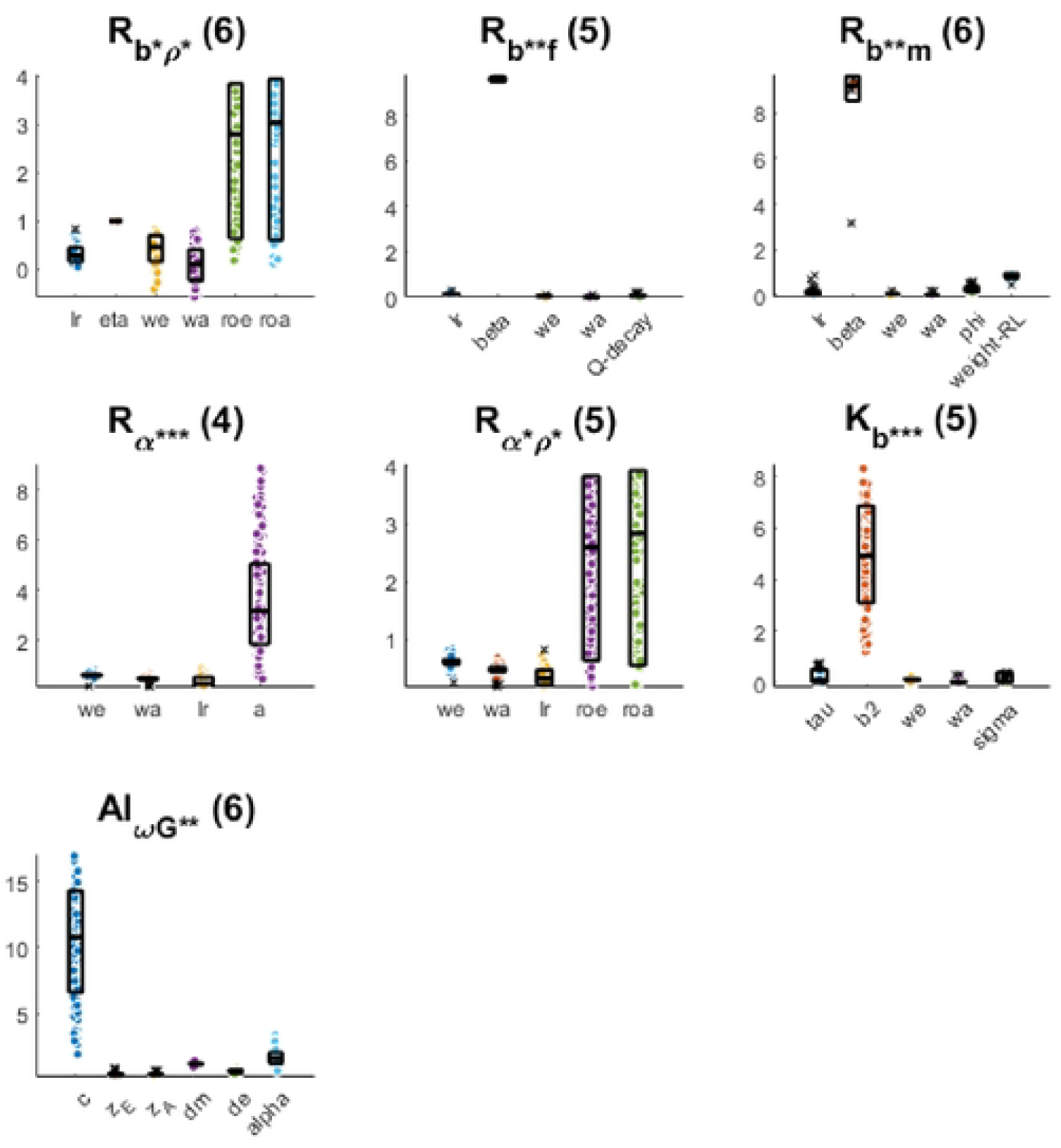
Fitted parameter ranges for individually hierarchically fitted models of the selected subspace for Dataset II.

**Figure S4.**
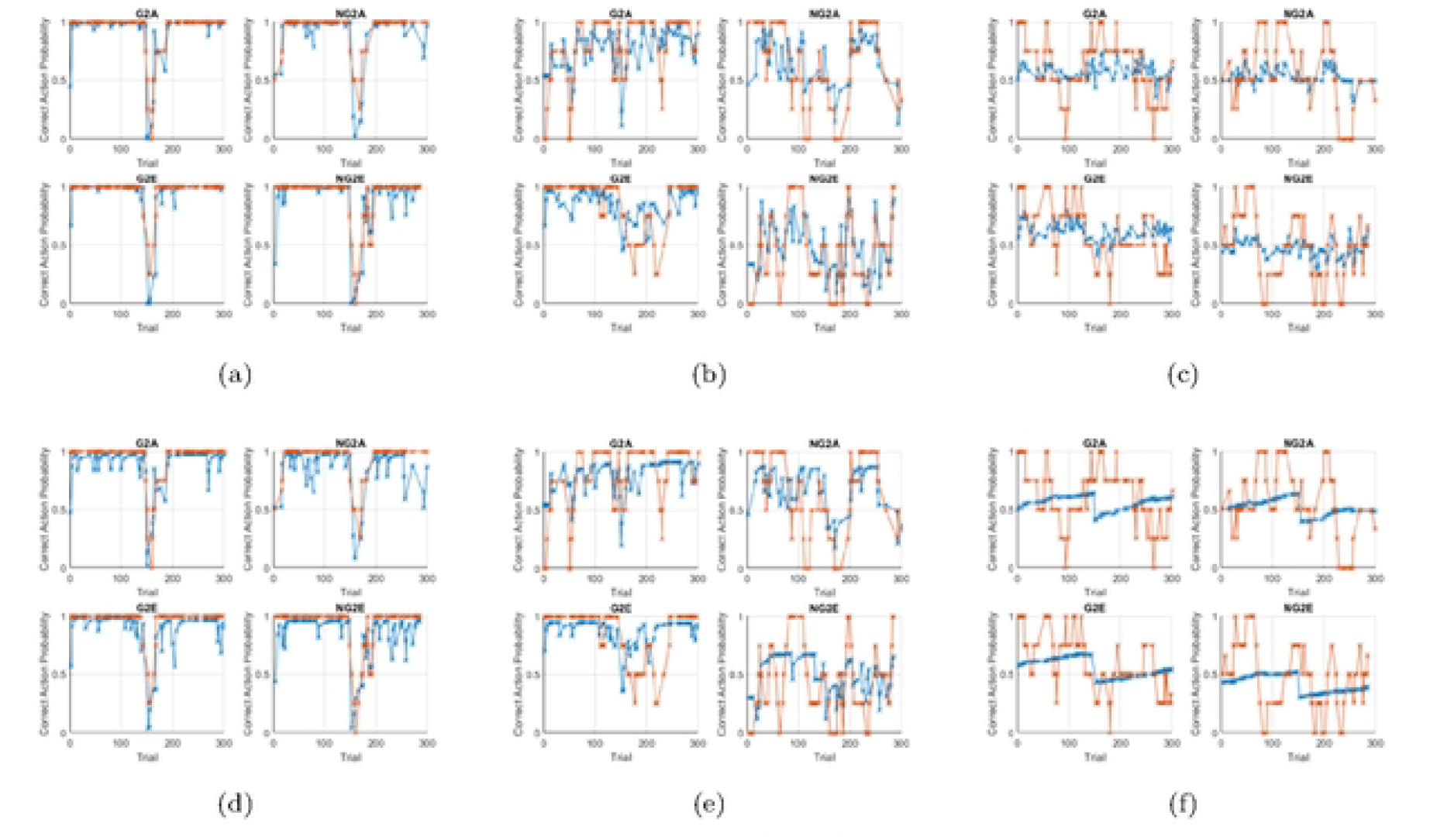
Exemplary model fits for Dataset I & II. (a)-(c) show the best, median and worst fit based on log-likelihood for the winning model for dataset I, (R_b**f_).(d)-(f) show the same subjects fitted by the worst performing model for dataset I, (A_ωG**_). Both models are hierarchically fitted to the full population. The plots show the observed choices (orange), smoothed with a rolling window of width four and the action probabilities based on the fitted model (blue) for each context.

**Figure S5.**
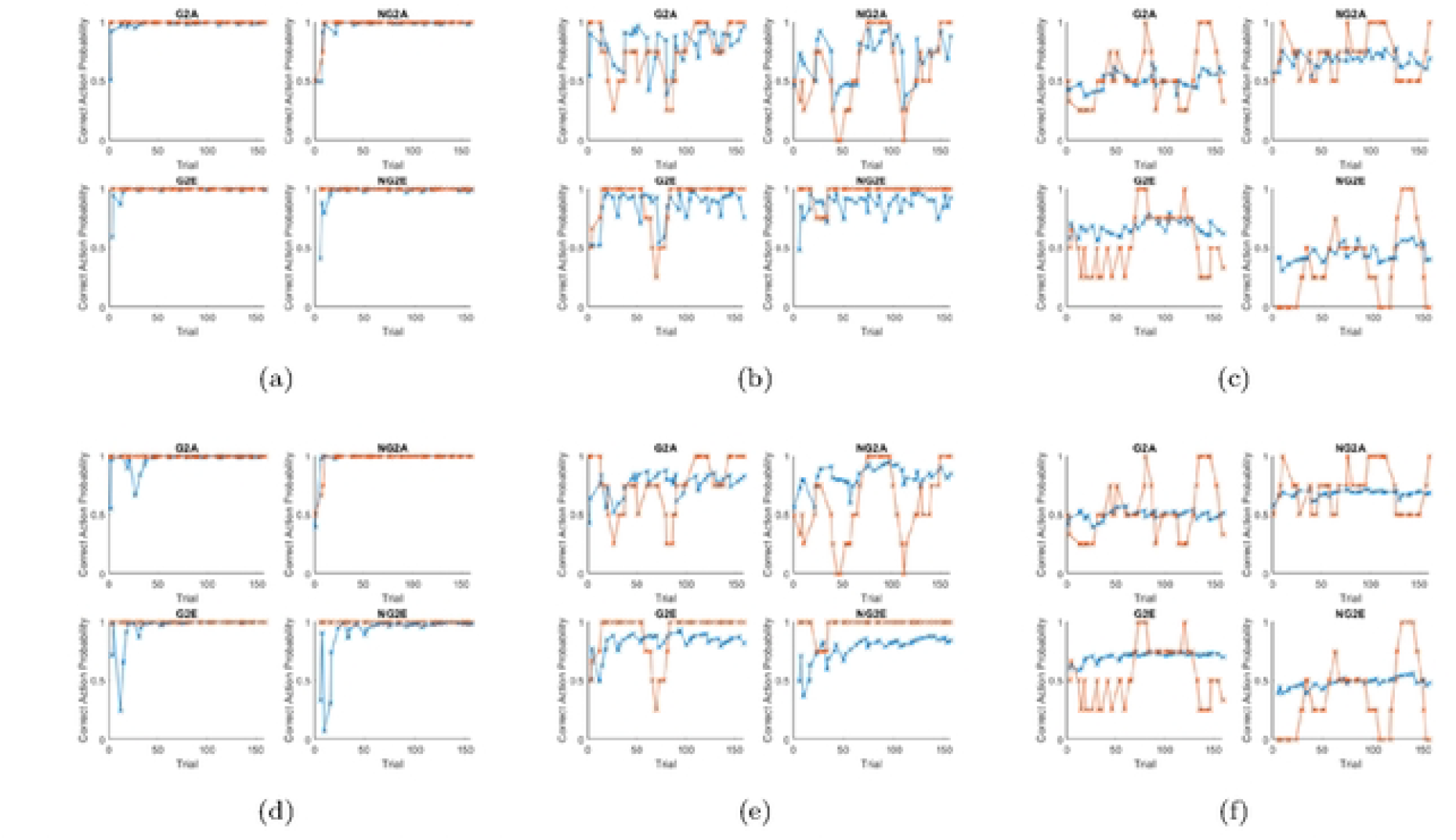
Exemplary model fits for Dataset II. (a)-(c) show the best, median and worst fit based on log-likelihood for the winning model for dataset II, (R_b**f_).(d)-(f) show the same subjects fitted by the worst performing model for dataset II, (A_ωG**_). Both models are hierarchically fitted to the full population. The plots show the observed choices (orange), smoothed with a rolling window of width four and the action probabilities based on the fitted model (blue) for each context.

**Figure S6.**
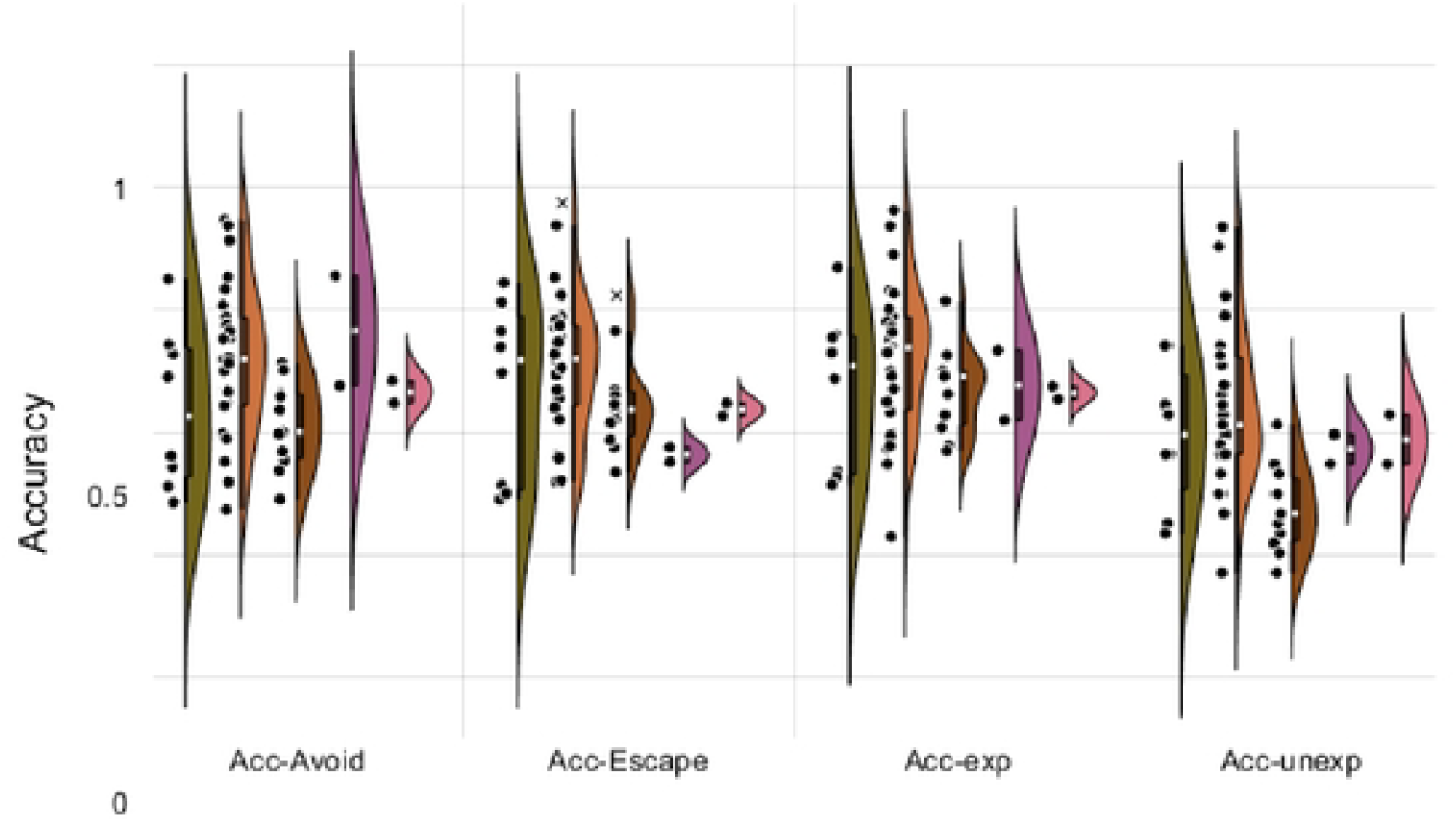
Subgroup Model-agnostic accuracy measures for Dataset I. *Subgroup Model-agnostic accuracy measures for Dataset I, including accuracy during avoid conditions, accuracy during escape conditions and accuracy on trials with expected feedback vs unexpected feedback. Significant differences between subgroups are indicated by star symbols (FDR-equivalent of \**: FDR_cuttoff = 4.4e-4 *- p<*.*05, **:* FDR_cuttoff = 0.0*- p<*.*01, ***:* FDR_cuttoff = 0.0*-p<*.*001). Significance is calculated according to the FDR-corrected p-value of a two-sided t-test. Significant differences between accuracy before and after reversal are uncorrected, as they are evaluated within each subgroup only*.

**Figure S7.**
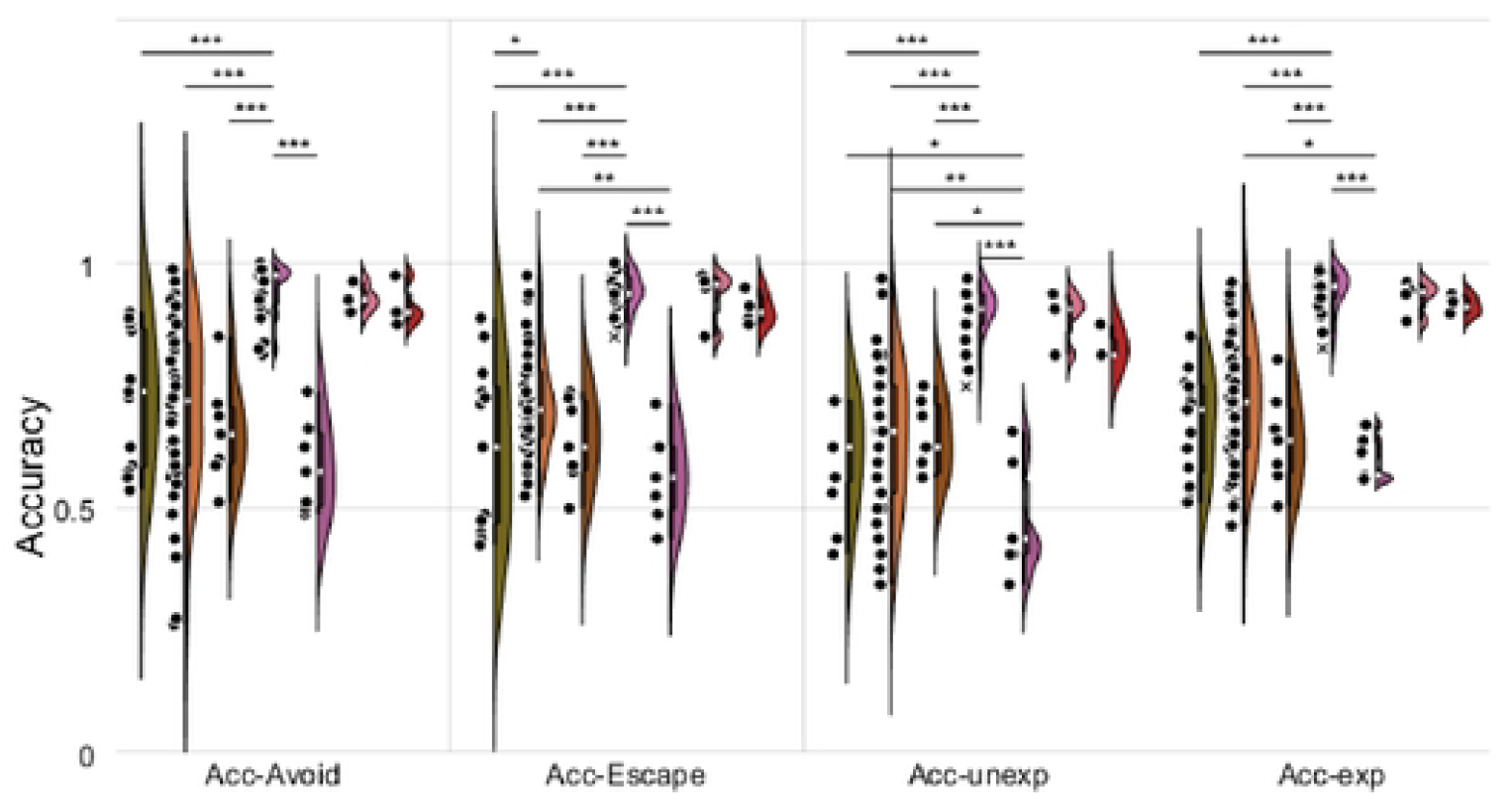
Subgroup Model-agnostic accuracy measures for Dataset II. Subgroup Model-agnostic accuracy measures for Dataset II, including accuracy during avoid conditions, accuracy during escape conditions and accuracy on trials with expected feedback vs unexpected feedback. Significant differences between subgroups are indicated by star symbols (FDR-equivalent of *: FDR_cuttoff = 4.4e-4 - p<.05, **: FDR_cuttoff = 0.0-p<.01, ***: FDR_cuttoff = 0.0-p<.001). Significance is calculated according to the FDR-corrected p-value of a two-sided t-test. Significant differences between accuracy before and after reversal are uncorrected, as they are evaluated within each subgroup only.

**Figure S8.**
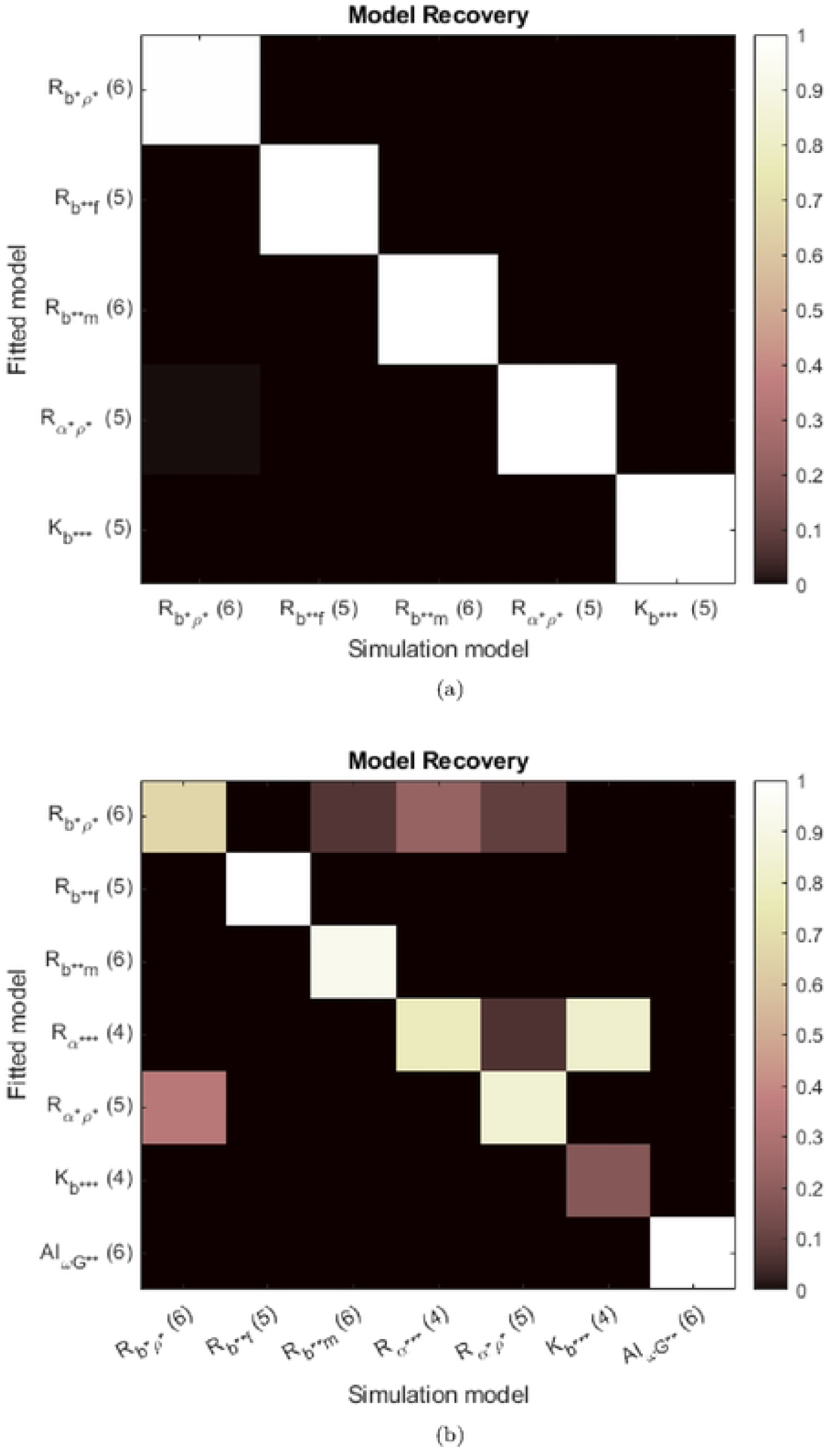
Model Recovery for Dataset I (a) and Dataset II (b) for the selected model subspace.

**Table S1.**
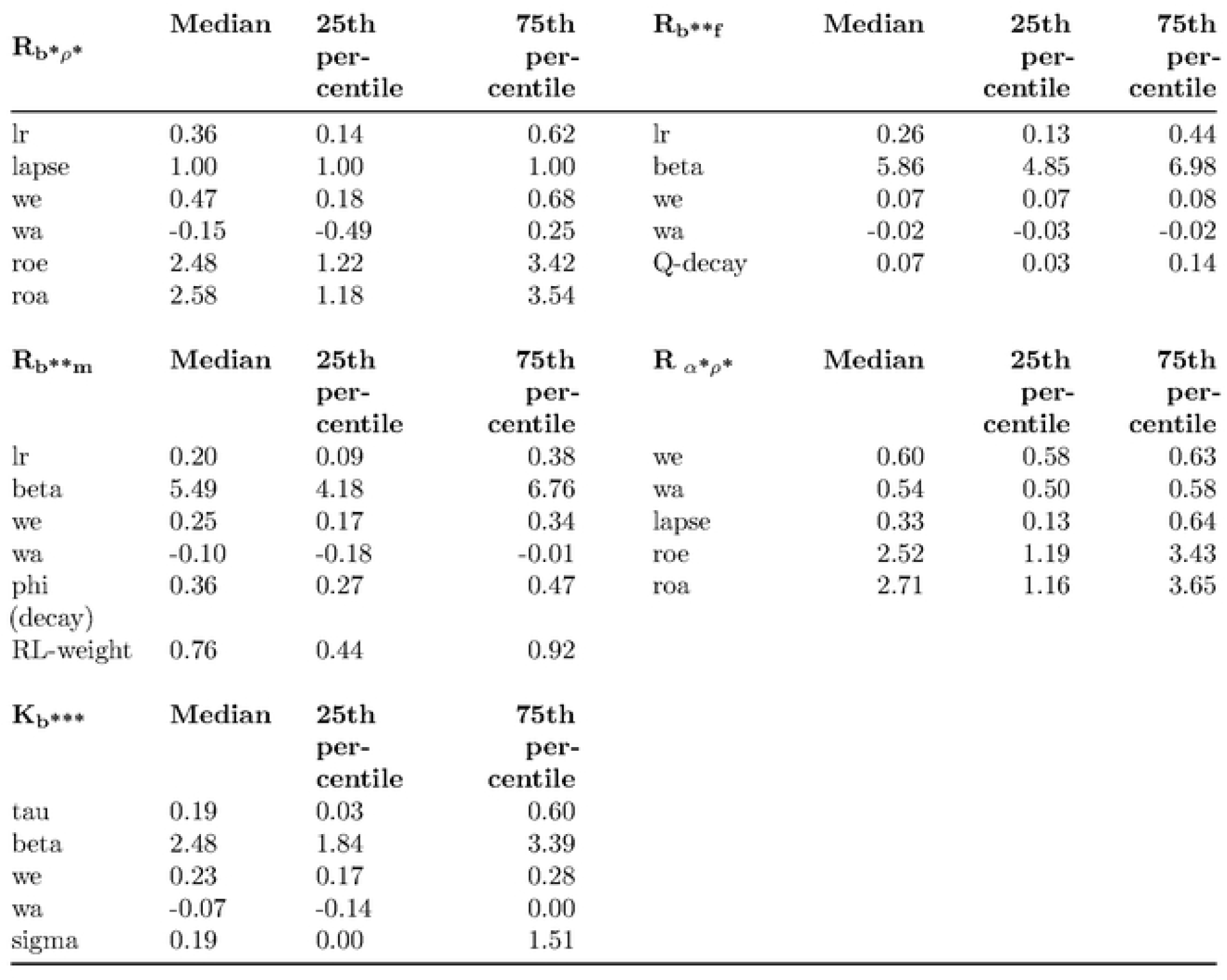
Distribution of fitted group prior for each model of the selected subspace for dataset I. Median, 25^th^ and 75% percentile are used to represent the skewness of prior distributions.

**Table S2.**
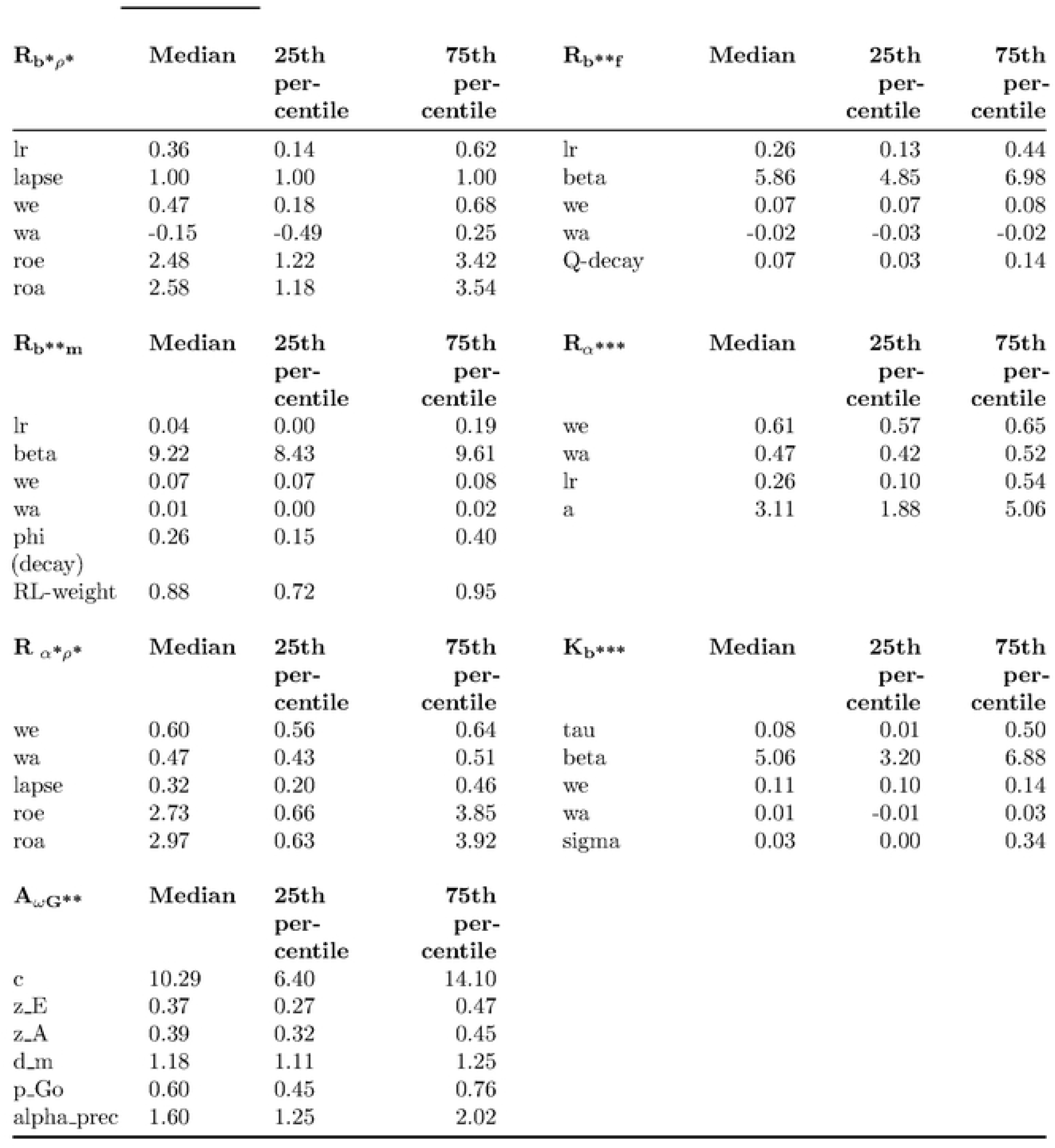
Distribution of fitted group prior for each model of the selected subspace for dataset II. Median, 25^th^ and 75% percentile are used to represent the skewness of prior distributions.

**Table S3.** Geometric average of predictive probability per choice, for datasets I and II, split by trial type. Best fitting model indicated in bold for each dataset and each trial type.

|  | Dataset I |  |  |  | Dataset II |  |  |  |
| --- | --- | --- | --- | --- | --- | --- | --- | --- |
|  | GA | NGA | GE | NGE | GA | NGA | GE | NGE |
| $R_{\omega G^{**}}$ (4) | 0.769 | 0.571 | 0.755 | 0.571 | 0.750 | 0.762 | 0.867 | 0.596 |
| $R_{\omega^* \rho^*}$ (5) | 0.740 | 0.576 | 0.759 | 0.573 | 0.762 | 0.706 | 0.817 | 0.642 |
| $R_{\nu G^{**}}$ (4) | 0.756 | 0.571 | 0.752 | 0.559 | 0.694 | 0.757 | 0.867 | 0.541 |
| $R_{\omega G^{**}}$ (3) | 0.754 | 0.584 | 0.748 | 0.581 | **0.775** | 0.746 | 0.806 | 0.695 |
| $R_{*G^{**}}$ (4) | 0.727 | 0.611 | 0.771 | 0.552 | 0.689 | 0.788 | 0.839 | 0.620 |
| $R_{b^* \rho^*}$ (6) | 0.720 | 0.616 | 0.761 | 0.565 | 0.711 | 0.766 | 0.816 | 0.639 |
| $R_{b^{**f}}$ (5) | 0.724 | 0.607 | 0.771 | 0.541 | 0.662 | 0.767 | 0.822 | 0.600 |
| $R_{b^{**m}}$ (6) | **0.788** | 0.580 | **0.860** | 0.464 | 0.646 | 0.785 | **0.892** | 0.510 |
| $R_{\alpha^{***}}$ (4) | 0.724 | 0.611 | 0.772 | 0.546 | 0.682 | 0.788 | 0.830 | 0.617 |
| $R_{\alpha^* \rho^*}$ (5) | 0.722 | 0.608 | 0.770 | 0.541 | 0.668 | 0.769 | 0.810 | 0.604 |
| $K_{b^{***}}$ (5) | 0.725 | 0.609 | 0.769 | 0.549 | 0.701 | **0.791** | 0.844 | 0.628 |
| $A_{\omega G^{**}}$ (6) | 0.699 | **0.636** | 0.724 | **0.627** | 0.719 | 0.745 | 0.797 | **0.675** |

**Table S4.** Model evidence for Datasets I & II. Logarithmic likelihood (loglik), logarithmic posterior (logpost), integrated log-likelihood (iL) and integrated BIC (iBIC) values for individually and hierarchically fitted candidate models.

|  | Dataset I |  |  |  |  |  |
| --- | --- | --- | --- | --- | --- | --- |
|  | Ind fit |  | Hier fit |  |  |  |
|  | ind. Loglik | ind.logpost | ind. Loglik | ind.logpost | iL | iBIC |
| $R_{\omega G^{**}}(4)$ | -142.41 | -150.75 | -142.65 | -148.24 | -148.08 | 297.70 |
| $R_{\omega^* \rho^*}(5)$ | -142.63 | -154.26 | -142.80 | -151.19 | -149.16 | 300.23 |
| $R_{\nu G^{**}}(4)$ | -149.41 | -158.58 | -149.42 | -155.76 | -155.69 | 312.91 |
| $R_{\omega G^{**}}(3)$ | -142.51 | -148.66 | -142.83 | -145.93 | -147.99 | 297.12 |
| $R_{*G^{**}}(4)$ | -139.58 | -147.52 | -140.09 | -143.94 | -146.79 | 295.12 |
| $R_{b^* \rho^*}(6)$ | -139.32 | -151.31 | -139.86 | -148.29 | -147.17 | 296.66 |
| $R_{b^{**f}}(5)$ | **-134.57** | **-144.53** | **-135.38** | **-137.71** | **-141.74** | **285.40** |
| $R_{b^{**m}}(6)$ | -135.20 | -146.79 | -136.22 | -142.81 | -144.44 | 291.19 |
| $R_{\alpha^{***}}(4)$ | -139.62 | -148.07 | -139.96 | -143.65 | -146.30 | 294.14 |
| $R_{\alpha^* \rho^*}(5)$ | -138.76 | -148.49 | -139.18 | -145.29 | -146.63 | 295.19 |
| $K_{b^{***}}(5)$ | -138.99 | -148.88 | -139.44 | -145.27 | -146.58 | 295.08 |
| $A_{\omega G^{**}}(6)$ | -143.07 | -154.18 | -144.03 | -150.75 | -175.83 | 353.97 |

|  | Dataset II |  |  |  |  |  |
| --- | --- | --- | --- | --- | --- | --- |
|  | Ind fit |  | Hier fit (1 model at a time) |  |  |  |
|  | ind. Loglik | ind.logpost | ind. Loglik | ind.logpost | iL | iBIC |
| $R_{\omega G^{**}}(4)$ | -62.22 | -70.79 | -62.45 | -68.23 | -67.09 | 134.87 |
| $R_{\omega^* \rho^*}(5)$ | -65.81 | -77.94 | -65.36 | -73.14 | -69.79 | 140.44 |
| $R_{\nu G^{**}}(4)$ | -69.26 | -78.27 | -68.69 | -75.24 | -73.54 | 147.77 |
| $R_{\omega G^{**}}(3)$ | -63.41 | -69.63 | -63.95 | -66.56 | -67.81 | 136.15 |
| $R_{*G^{**}}(4)$ | -60.64 | -68.63 | -61.35 | -64.43 | -66.36 | 133.40 |
| $R_{b^* \rho^*}(6)$ | -61.97 | -74.02 | -62.32 | -70.54 | -67.43 | 135.89 |
| $R_{b^{**f}}(5)$ | **-59.19** | -69.13 | **-60.26** | **-60.16** | **-64.15** | **129.16** |
| $R_{b^{**m}}(6)$ | -59.51 | -71.39 | -60.54 | -64.89 | -65.80 | 132.63 |
| $R_{\alpha^{***}}(4)$ | -60.36 | **-68.74** | -60.98 | -65.10 | -65.97 | 132.62 |
| $R_{\alpha^* \rho^*}(5)$ | -61.11 | -70.98 | -61.65 | -67.97 | -67.42 | 135.70 |
| $K_{b^{***}}(5)$ | -60.29 | -70.27 | -60.92 | -65.72 | -66.17 | 133.20 |
| $A_{\omega G^{**}}(6)$ | -60.78 | -72.12 | -61.77 | -67.45 | -85.19 | 171.41 |

**Table S5.** p-values for subgrouping analysis of Model-agnostic behavioural measures for Dataset I, including performance biases (overall Go-Bias, Excitatory-Escape Bias and Inhibitory-Avoid Bias), learning strategies (Lose-Switch and Win-Stay behaviours) and accuracy (total, Before and After reversal). Significant differences between subgroups are indicated by star symbols (FDR-equivalent of * - p*<*.05, **- p*<*.01, ***-p*<*.001). Significance is calculated according to the FDR-corrected p-value of a two-sided t-test.

|  | thresholds | 0.05 | 0.01 | 0.001 |  |
| --- | --- | --- | --- | --- | --- |
|  | FDR | 4.40E-04 | 0 | 0 |  |
| **GE** |  |  |  |  |  |
| | $R_{b^*\rho^*}$ (6) | $R_{b^{**f}}$ (5) | $R_{b^{**m}}$ (6) | $R_{\alpha^*\rho^*}$ (5) | $K_{b^{***}}$ (5) |
| $R_{b^*\rho^*}$ (6) | | 0.31797 | 0.241676 | 0.382566 | 0.466726 |
| $R_{b^{**f}}$ (5) | | | 0.924886 | 2.49E-01 | 0.920912 |
| $R_{b^{**m}}$ (6) | | | | 1.27E-01 | 0.852999 |
| $R_{\alpha^*\rho^*}$ (5) | | | | | 0.505305 |
| $K_{b^{***}}$ (5) | | | | | |
| **NGA** |  |  |  |  |  |
| | $R_{b^*\rho^*}$ (6) | $R_{b^{**f}}$ (5) | $R_{b^{**m}}$ (6) | $R_{\alpha^*\rho^*}$ (5) | $K_{b^{***}}$ (5) |
| $R_{b^*\rho^*}$ (6) | | 0.048671 | 6.62E-02 | 6.42E-03 | 6.74E-02 |
| $R_{b^{**f}}$ (5) | | | 0.679877 | 0.093524 | 8.13E-01 |
| $R_{b^{**m}}$ (6) | | | | 0.059215 | 6.23E-01 |
| $R_{\alpha^*\rho^*}$ (5) | | | | | 4.93E-01 |
| $K_{b^{***}}$ (5) | | | | | |
| **Go** |  |  |  |  |  |
| | $R_{b^*\rho^*}$ (6) | $R_{b^{**f}}$ (5) | $R_{b^{**m}}$ (6) | $R_{\alpha^*\rho^*}$ (5) | $K_{b^{***}}$ (5) |
| $R_{b^*\rho^*}$ (6) | | 1.08E-01 | 5.20E-02 | 0.083634 | 0.202065 |
| $R_{b^{**f}}$ (5) | | | 0.800619 | 8.25E-01 | 8.66E-01 |
| $R_{b^{**m}}$ (6) | | | | 5.80E-01 | 0.695301 |
| $R_{\alpha^*\rho^*}$ (5) | | | | | 9.76E-01 |
| $K_{b^{***}}$ (5) | | | | | |
| **Lose-Switch** |  |  |  |  |  |
| | $R_{b^*\rho^*}$ (6) | $R_{b^{**f}}$ (5) | $R_{b^{**m}}$ (6) | $R_{\alpha^*\rho^*}$ (5) | $K_{b^{***}}$ (5) |
| $R_{b^*\rho^*}$ (6) | | 0.047858 | 0.001172 | 4.75E-01 | 0.000963 |
| $R_{b^{**f}}$ (5) | | | 1.68E-05 | 5.50E-01 | 0.250171 |
| $R_{b^{**m}}$ (6) | | | | 2.30E-02 | 4.19E-05 |
| $R_{\alpha^*\rho^*}$ (5) | | | | | 0.230332 |
| $K_{b^{***}}$ (5) | | | | | |
| **Win-Stay** |  |  |  |  |  |
| | $R_{b^*\rho^*}$ (6) | $R_{b^{**f}}$ (5) | $R_{b^{**m}}$ (6) | $R_{\alpha^*\rho^*}$ (5) | $K_{b^{***}}$ (5) |
| $R_{b^*\rho^*}$ (6) | | 0.041767 | 0.931062 | 7.85E-01 | 0.222764 |
| $R_{b^{**f}}$ (5) | | | 0.012535 | 3.90E-01 | 0.665042 |
| $R_{b^{**m}}$ (6) | | | | 6.63E-01 | 0.03233 |
| $R_{\alpha^*\rho^*}$ (5) | | | | | 2.35E-01 |
| $K_{b^{***}}$ (5) | | | | | |
| **Accuracy** |  |  |  |  |  |
| | $R_{b^*\rho^*}$ (6) | $R_{b^{**f}}$ (5) | $R_{b^{**m}}$ (6) | $R_{\alpha^*\rho^*}$ (5) | $K_{b^{***}}$ (5) |
| $R_{b^*\rho^*}$ (6) | | 0.216057 | 0.581143 | 9.19E-01 | 0.972961 |
| $R_{b^{**f}}$ (5) | | | 0.027575 | 5.54E-01 | 0.45129 |
| $R_{b^{**m}}$ (6) | | | | 4.95E-01 | 0.660695 |
| $R_{\alpha^*\rho^*}$ (5) | | | | | 7.79E-01 |
| $K_{b^{***}}$ (5) | | | | | |

| **Accuracy-**<br>**Before** |  |  |  |  |  |
| --- | --- | --- | --- | --- | --- |
| | $R_{b^*\rho^*}$ (6) | $R_{b^{**f}}$ (5) | $R_{b^{**m}}$ (6) | $R_{\alpha^*\rho^*}$ (5) | $K_{b^{***}}$ (5) |
| $R_{b^*\rho^*}$ (6) | | 0.476048 | 0.100235 | 5.97E-01 | 0.084144 |
| $R_{b^{**f}}$ (5) | | | 0.006269 | 3.38E-01 | 0.11522 |
| $R_{b^{**m}}$ (6) | | | | 6.29E-01 | 0.001241 |
| $R_{\alpha^*\rho^*}$ (5) | | | | | 3.17E-02 |
| $K_{b^{***}}$ (5) | | | | | |
| **Accuracy-**<br>**After** |  |  |  |  |  |
| | $R_{b^*\rho^*}$ (6) | $R_{b^{**f}}$ (5) | $R_{b^{**m}}$ (6) | $R_{\alpha^*\rho^*}$ (5) | $K_{b^{***}}$ (5) |
| $R_{b^*\rho^*}$ (6) | | 0.180973 | 0.37986 | 5.06E-01 | 0.093958 |
| $R_{b^{**f}}$ (5) | | | 0.52155 | 9.54E-01 | 0.004086 |
| $R_{b^{**m}}$ (6) | | | | 5.53E-01 | 0.000409 |
| $R_{\alpha^*\rho^*}$ (5) | | | | | 1.90E-02 |
| $K_{b^{***}}$ (5) | | | | | |

**Table S6.** p-values for subgrouping analysis of Model-agnostic behavioural measures for Dataset II, including performance biases (overall Go-Bias, Excitatory-Escape Bias and Inhibitory-Avoid Bias), learning strategies (Lose-Switch and Win-Stay behaviours) and accuracy (total, Escape and Avoid condition). Significant differences between subgroups are indicated by star symbols (FDR-equivalent of * - p*<*.05, **- p*<*.01, ***-p*<*.001). Significance is calculated according to the FDR-corrected p-value of a two-sided t-test.

|  | thresholds<br>FDR | 0.05<br>0.021 | 0.01<br>0.003 | 0.001<br>0.0002 |  |  |  |
| --- | --- | --- | --- | --- | --- | --- | --- |
| **GE** |  |  |  |  |  |  |  |
| | $R_{b^*\rho^*}$ (6) | $R_{b^{**f}}$ (5) | $R_{b^{**m}}$ (6) | $R_{\alpha^{***}}$ (4) | $R_{\alpha^*\rho^*}$ (5) | $K_{b^{***}}$ (5) | $A_{\omega G^{**}}$ (6) |
| $R_{b^*\rho^*}$ (6) | 0 | 0.000595 | 0.0259 | 0.190799 | 0.815467 | 0.827384 | 0.875949 |
| $R_{b^{**f}}$ (5) | | 0 | 0.885315 | 1.68E-09 | 0.1216 | 0.021138 | 0.024241 |
| $R_{b^{**m}}$ (6) | | | 0 | 1.04E-06 | 0.521065 | 0.052075 | 0.055171 |
| $R_{\alpha^{***}}$ (4) | | | | 0 | 0.429624 | 0.404827 | 0.310596 |
| $R_{\alpha^*\rho^*}$ (5) | | | | | 0 | 0.847083 | 0.861032 |
| $K_{b^{***}}$ (5) | | | | | | 0 | 0.915152 |
| $A_{\omega G^{**}}$ (6) | | | | | | | 0 |
| **NGA** |  |  |  |  |  |  |  |
| | $R_{b^*\rho^*}$ (6) | $R_{b^{**f}}$ (5) | $R_{b^{**m}}$ (6) | $R_{\alpha^{***}}$ (4) | $R_{\alpha^*\rho^*}$ (5) | $K_{b^{***}}$ (5) | $A_{\omega G^{**}}$ (6) |
| $R_{b^*\rho^*}$ (6) | | 0.000155 | 5.79E-05 | 1.50E-08 | 1.67E-05 | 0.000292 | 0.054584 |
| $R_{b^{**f}}$ (5) | | | 0.096034 | 0.918499 | 4.36E-11 | 0.542572 | 0.326606 |
| $R_{b^{**m}}$ (6) | | | | 0.005857 | 8.89E-06 | 0.486716 | 0.048786 |
| $R_{\alpha^{***}}$ (4) | | | | | 1.11E-14 | 0.120472 | 0.02991 |
| $R_{\alpha^*\rho^*}$ (5) | | | | | | 8.32E-05 | 0.000592 |
| $K_{b^{***}}$ (5) | | | | | | | 0.005319 |
| $A_{\omega G^{**}}$ (6) | | | | | | | |
| **Go** |  |  |  |  |  |  |  |
| | $R_{b^*\rho^*}$ (6) | $R_{b^{**f}}$ (5) | $R_{b^{**m}}$ (6) | $R_{\alpha^{***}}$ (4) | $R_{\alpha^*\rho^*}$ (5) | $K_{b^{***}}$ (5) | $A_{\omega G^{**}}$ (6) |
| $R_{b^*\rho^*}$ (6) | | 9.96E-07 | 2.67E-05 | 0.000415 | 0.04157 | 0.009729 | 0.218667 |
| $R_{b^{**f}}$ (5) | | | 0.266029 | 1.74E-05 | 5.15E-08 | 0.252848 | 0.027146 |
| $R_{b^{**m}}$ (6) | | | | 3.41E-06 | 0.002378 | 0.079696 | 0.013927 |
| $R_{\alpha^{***}}$ (4) | | | | | 3.62E-05 | 0.139608 | 0.327132 |
| $R_{\alpha^*\rho^*}$ (5) | | | | | | 0.058708 | 0.133694 |
| $K_{b^{***}}$ (5) | | | | | | | 0.054934 |
| $A_{\omega G^{**}}$ (6) | | | | | | | |
| **Lose-Switch** |  |  |  |  |  |  |  |
| | $R_{b^*\rho^*}$ (6) | $R_{b^{**f}}$ (5) | $R_{b^{**m}}$ (6) | $R_{\alpha^{***}}$ (4) | $R_{\alpha^*\rho^*}$ (5) | $K_{b^{***}}$ (5) | $A_{\omega G^{**}}$ (6) |
| $R_{b^*\rho^*}$ (6) | | 0.026847 | 0.295972 | 1.09E-10 | 0.00121 | 0.010088 | 0.008872 |
| $R_{b^{**f}}$ (5) | | | 0.003107 | 2.88E-14 | 0.003254 | 0.032104 | 0.049205 |
| $R_{b^{**m}}$ (6) | | | | 2.63E-10 | 0.000411 | 0.003035 | 0.00088 |
| $R_{\alpha^{***}}$ (4) | | | | | 0.033357 | 0.059416 | 0.024521 |
| $R_{\alpha^*\rho^*}$ (5) | | | | | | 0.896589 | 0.72132 |
| $K_{b^{***}}$ (5) | | | | | | | 0.855777 |
| $A_{\omega G^{**}}$ (6) | | | | | | | |
| **Win-Stay** |  |  |  |  |  |  |  |
| | $R_{b^*\rho^*}$ (6) | $R_{b^{**f}}$ (5) | $R_{b^{**m}}$ (6) | $R_{\alpha^{***}}$ (4) | $R_{\alpha^*\rho^*}$ (5) | $K_{b^{***}}$ (5) | $A_{\omega G^{**}}$ (6) |
| $R_{b^*\rho^*}$ (6) | | 0.000442 | 0.74775 | 2.01E-16 | 0.065452 | 4.90E-05 | 9.63E-05 |
| $R_{b^{**f}}$ (5) | | | 0.020246 | 3.74E-14 | 0.512338 | 0.000275 | 0.000908 |
| $R_{b^{**m}}$ (6) | | | | 1.15E-12 | 0.21552 | 0.001467 | 0.002309 |
| $R_{\alpha^{***}}$ (4) | | | | | 3.50E-14 | 0.16952 | 0.752648 |
| $R_{\alpha^*\rho^*}$ (5) | | | | | | 0.00028 | 0.000581 |
| K<sub>b***</sub> (5) |  |  |  |  |  |  | 0.126631 |
| A<sub>ωG**</sub> (6) |  |  |  |  |  |  |  |
| **Accuracy** |  |  |  |  |  |  |  |
|  | R<sub>b*ρ*</sub> (6) | R<sub>b**f</sub> (5) | R<sub>b**m</sub> (6) | R<sub>α***</sub> (4) | R<sub>α*ρ*</sub> (5) | K<sub>b***</sub> (5) | A<sub>ωG**</sub> (6) |
| R<sub>b*ρ*</sub> (6) |  | 0.271798 | 0.604591 | 2.74E-14 | 0.04016 | 0.000239 | 0.000476 |
| R<sub>b**f</sub> (5) |  |  | 0.17584 | 6.95E-15 | 0.005399 | 0.00068 | 0.001964 |
| R<sub>b**m</sub> (6) |  |  |  | 1.12E-14 | 0.084754 | 0.000132 | 0.000181 |
| R<sub>α***</sub> (4) |  |  |  |  | 1.75E-19 | 0.611789 | 0.130608 |
| R<sub>α*ρ*</sub> (5) |  |  |  |  |  | 8.21E-07 | 6.59E-07 |
| K<sub>b***</sub> (5) |  |  |  |  |  |  | 0.336925 |
| A<sub>ωG**</sub> (6) |  |  |  |  |  |  |  |
| **Accuracy-Avoid** |  |  |  |  |  |  |  |
|  | R<sub>b*ρ*</sub> (6) | R<sub>b**f</sub> (5) | R<sub>b**m</sub> (6) | R<sub>α***</sub> (4) | R<sub>α*ρ*</sub> (5) | K<sub>b***</sub> (5) | A<sub>ωG**</sub> (6) |
| R<sub>b*ρ*</sub> (6) |  | 0.740636 | 0.290712 | 3.79E-09 | 0.028745 | 0.008544 | 0.014229 |
| R<sub>b**f</sub> (5) |  |  | 0.487101 | 5.93E-10 | 0.081488 | 0.012723 | 0.020137 |
| R<sub>b**m</sub> (6) |  |  |  | 2.16E-11 | 0.211863 | 0.00094 | 0.001631 |
| R<sub>α***</sub> (4) |  |  |  |  | 1.36E-14 | 0.54201 | 0.259552 |
| R<sub>α*ρ*</sub> (5) |  |  |  |  |  | 6.38E-05 | 0.000116 |
| K<sub>b***</sub> (5) |  |  |  |  |  |  | 0.557947 |
| A<sub>ωG**</sub> (6) |  |  |  |  |  |  |  |
| **Accuracy-Escape** |  |  |  |  |  |  |  |
|  | R<sub>b*ρ*</sub> (6) | R<sub>b**f</sub> (5) | R<sub>b**m</sub> (6) | R<sub>α***</sub> (4) | R<sub>α*ρ*</sub> (5) | K<sub>b***</sub> (5) | A<sub>ωG**</sub> (6) |
| R<sub>b*ρ*</sub> (6) |  | 0.012706 | 0.824285 | 1.53E-11 | 0.460558 | 0.002683 | 0.004871 |
| R<sub>b**f</sub> (5) |  |  | 0.062341 | 8.17E-16 | 0.001065 | 0.000297 | 0.001341 |
| R<sub>b**m</sub> (6) |  |  |  | 4.37E-15 | 0.189121 | 0.000203 | 0.000268 |
| R<sub>α***</sub> (4) |  |  |  |  | 3.76E-17 | 0.805182 | 0.109418 |
| R<sub>α*ρ*</sub> (5) |  |  |  |  |  | 6.23E-05 | 7.55E-05 |
| K<sub>b***</sub> (5) |  |  |  |  |  |  | 0.427067 |
| A<sub>ωG**</sub> (6) |  |  |  |  |  |  |  |

**Table S7.**
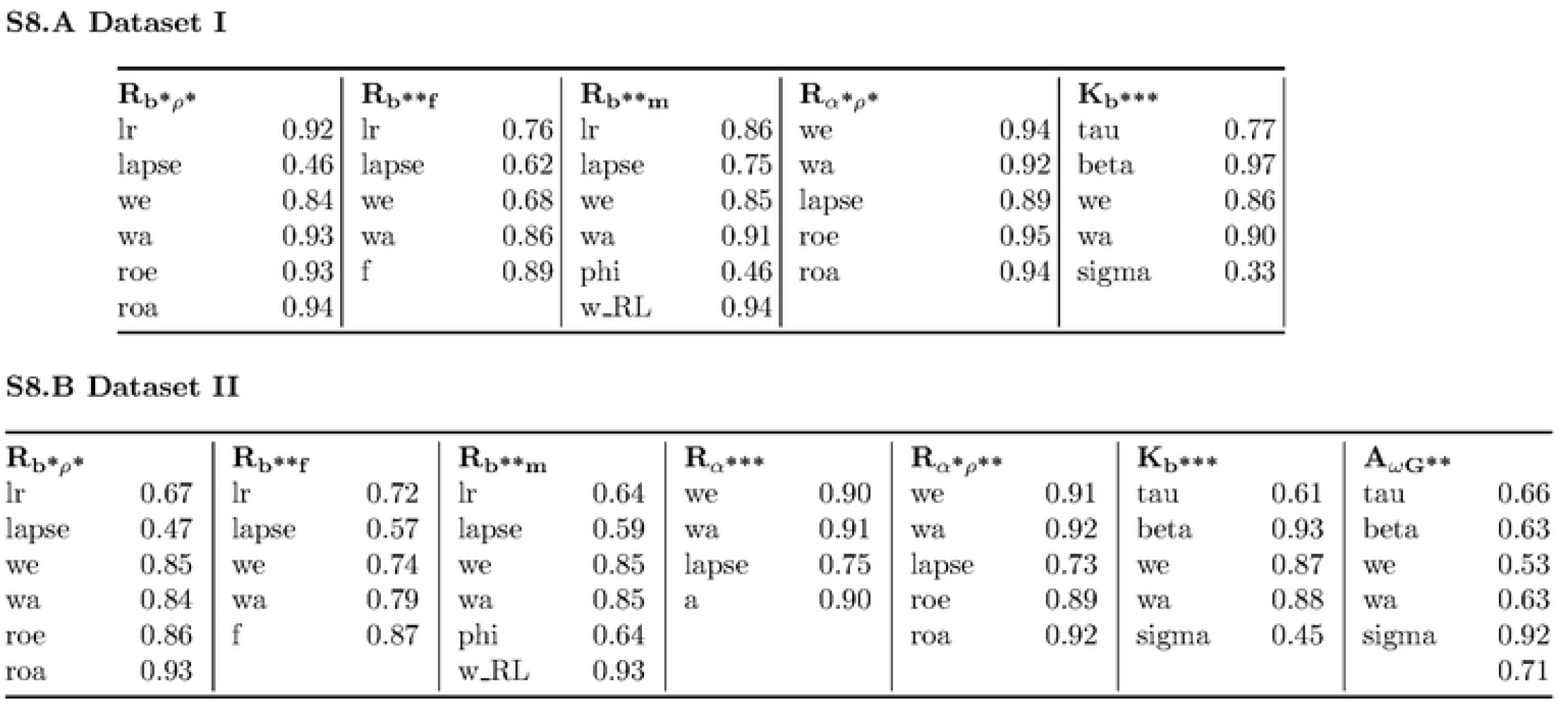
ICC values for parameter recovery for all models of the pre-selected subspace for datasets I &II, based on 50 and 114 simulated trajectories, respectively.

| $\mathbf{R}_{b^*\rho^*}$ | | $\mathbf{R}_{b^{**}f}$ | | $\mathbf{R}_{b^{**}m}$ | | $\mathbf{R}_{\alpha^*\rho^*}$ | | $\mathbf{K}_{b^{***}}$ | |
| --- | --- | --- | --- | --- | --- | --- | --- | --- | --- |
| lr | 0.92 | lr | 0.76 | lr | 0.86 | we | 0.94 | tau | 0.77 |
| lapse | 0.46 | lapse | 0.62 | lapse | 0.75 | wa | 0.92 | beta | 0.97 |
| we | 0.84 | we | 0.68 | we | 0.85 | lapse | 0.89 | we | 0.86 |
| wa | 0.93 | wa | 0.86 | wa | 0.91 | roe | 0.95 | wa | 0.90 |
| roe | 0.93 | f | 0.89 | phi | 0.46 | roa | 0.94 | sigma | 0.33 |
| roa | 0.94 |  |  | w_RL | 0.94 |  |  |  |  |

| $\mathbf{R}_{b^*\rho^*}$ | | $\mathbf{R}_{b^{**}f}$ | | $\mathbf{R}_{b^{**}m}$ | | $\mathbf{R}_{\alpha^{***}}$ | | $\mathbf{R}_{\alpha^*\rho^{**}}$ | | $\mathbf{K}_{b^{***}}$ | | $\mathbf{A}_{\omega G^{**}}$ | |
| --- | --- | --- | --- | --- | --- | --- | --- | --- | --- | --- | --- | --- | --- |
| lr | 0.67 | lr | 0.72 | lr | 0.64 | we | 0.90 | we | 0.91 | tau | 0.61 | tau | 0.66 |
| lapse | 0.47 | lapse | 0.57 | lapse | 0.59 | wa | 0.91 | wa | 0.92 | beta | 0.93 | beta | 0.63 |
| we | 0.85 | we | 0.74 | we | 0.85 | lapse | 0.75 | lapse | 0.73 | we | 0.87 | we | 0.53 |
| wa | 0.84 | wa | 0.79 | wa | 0.85 | a | 0.90 | roe | 0.89 | wa | 0.88 | wa | 0.63 |
| roe | 0.86 | f | 0.87 | phi | 0.64 |  |  | roa | 0.92 | sigma | 0.45 | sigma | 0.92 |
| roa | 0.93 |  |  | w_RL | 0.93 |  |  |  |  |  |  |  | 0.71 |

**Table S8.** Dataset I&II: ICC values indicting each model’s ability to replicate behavioural measures in simulated data based on a hierarchical fit of the model to the entire population.

Dataset I
| ICC Values | GE | NGA | Lswitch | Wstay | Accuracy before | Accuracy after |
| --- | --- | --- | --- | --- | --- | --- |
| $R_{\omega G^{**}}$ | 0.690 | 0.577 | 0.631 | 0.794 | 0.775 | 0.784 |
| $R_{\omega^* \rho^*}$ | 0.717 | 0.656 | 0.361 | 0.653 | 0.752 | 0.823 |
| $R_{\nu G^{**}}$ | 0.701 | 0.688 | 0.646 | 0.798 | 0.720 | 0.662 |
| $R^* G^{**}$ | 0.724 | 0.752 | 0.648 | 0.762 | 0.751 | 0.823 |
| $R_b^{***}$ | **0.949** | 0.957 | 0.656 | 0.791 | 0.791 | **0.843** |
| $R_b^* \rho^*$ | 0.890 | 0.929 | 0.385 | 0.721 | 0.733 | 0.796 |
| $R_b^{**f}$ | 0.913 | 0.947 | 0.662 | **0.908** | 0.840 | 0.735 |
| $R_b^{**m}$ | 0.741 | 0.885 | **0.734** | 0.837 | 0.498 | 0.712 |
| $R_{\alpha}^{***}$ | 0.918 | 0.946 | 0.629 | 0.787 | 0.826 | 0.804 |
| $R_{\alpha^* \rho^*}$ | 0.944 | **0.962** | 0.297 | 0.739 | 0.747 | 0.816 |
| $K_b^{***}$ | 0.891 | 0.931 | 0.651 | 0.830 | 0.772 | 0.734 |
| $A_{\omega G^{**}}$ | 0.712 | 0.788 | 0.814 | 0.785 | **0.846** | 0.723 |

| ICC Values | GE | NGA | Lswitch | Wstay | Accuracy |
| --- | --- | --- | --- | --- | --- |
| $R_{\omega G^{**}}$ | 0.762 | 0.665 | 0.751 | 0.859 | 0.902 |
| $R_{\omega^* \rho^*}$ | 0.428 | 0.141 | 0.444 | 0.794 | 0.843 |
| $R_{\nu G^{**}}$ | 0.573 | 0.587 | 0.584 | 0.761 | 0.828 |
| $R^* G^{**}$ | 0.532 | 0.534 | 0.717 | 0.853 | 0.903 |
| $R_b^{***}$ | 0.862 | 0.934 | 0.834 | 0.864 | 0.936 |
| $R_b^* \rho^*$ | 0.770 | 0.756 | 0.412 | 0.797 | 0.874 |
| $R_b^{**f}$ | 0.865 | 0.892 | 0.700 | **0.901** | 0.923 |
| $R_b^{**m}$ | 0.889 | 0.923 | 0.765 | 0.852 | 0.910 |
| $R_{\alpha}^{***}$ | 0.912 | 0.925 | 0.738 | 0.865 | 0.953 |
| $R_{\alpha^* \rho^*}$ | **0.918** | 0.930 | 0.485 | 0.786 | 0.908 |
| $K_b^{***}$ | 0.909 | 0.924 | 0.790 | 0.854 | 0.932 |
| $A_{\omega G^{**}}$ | 0.904 | **0.945** | **0.837** | 0.823 | **0.973** |

